# Robust differential composition and variability analysis for multisample cell omics

**DOI:** 10.1101/2022.03.04.482758

**Authors:** S Mangiola, A Schulze, M Trussart, E Zozaya, M Ma, Z Gao, AF Rubin, TP Speed, H Shim, AT Papenfuss

**Affiliations:** The Walter and Eliza Hall Institute of Medical Research, Parkville, Victoria, Australia; Department of Medical Biology, University of Melbourne, Melbourne, Victoria, Australia; Melbourne Integrative Genomics / School of Mathematics and Statistics, University of Melbourne, Victoria, Australia; Peter MacCallum Cancer Centre, Melbourne, VIC 3000, Australia; Sir Peter MacCallum Department of Oncology, University of Melbourne, Melbourne, Victoria, Australia

**Keywords:** single-cell RNA sequencing, CyTOF, microbiome, composition analysis, Bayes

## Abstract

Cell omics such as single-cell genomics, proteomics and microbiomics allow the characterisation of tissue and microbial community composition, which can be compared between conditions to identify biological drivers. This strategy has been critical to unveiling markers of disease progression such as cancer and pathogen infection. For cell omic data, no method for differential variability analysis exists, and methods for differential composition analysis only take a few fundamental data properties into account. Here we introduce sccomp, a generalised method for differential composition and variability analyses able to jointly model data count distribution, compositionality, group-specific variability and proportion mean-variability association, with awareness against outliers. Sccomp is an extensive analysis framework that allows realistic data simulation and cross-study knowledge transfer. Here, we demonstrate that mean-variability association is ubiquitous across technologies showing the inadequacy of the very popular Dirichlet-multinomial modelling and provide mandatory principles for differential variability analysis. We show that sccomp accurately fits experimental data, with a 50% incremental improvement over state-of-the-art algorithms. Using sccomp, we identified novel differential constraints and composition in the microenvironment of primary breast cancer.

**Significance statement:** Determining the composition of cell populations is made possible by technologies like single-cell transcriptomics, CyTOF and microbiome sequencing. Such analyses are now widespread across fields (~800 publications/month, Scopus). However, existing methods for differential abundance do not model all data features, and cell-type/taxa specific differential variability is not yet possible. Increase in the variability of tissue composition and microbial communities is a well-known indicator of loss of homeostasis and disease. A suitable statistical method would enable new types of analyses to identify component-specific loss of homeostasis for the first time. This and other innovations are now possible through our discovery of the mean-variability association for compositional data. Based on this fundamental observation, we have developed a new statistical model, sccomp, that enables differential variability analysis for composition data, improved differential abundance analyses, with cross-sample information borrowing, outlier identification and exclusion, realistic data simulation, based on experimental datasets, cross-study knowledge transfer.

## Introduction

Compositional analyses are central in many fields of biology. Tissue composition analysis enabled seminal discoveries in cancer research (1–6) and epidemiology (e.g. COVID19 (4)). Compositional analysis of microbial communities is crucial for studying metabolic disease (7) and skin physiology (8). Single-cell transcriptomics (9) and high-throughput flow cytometry (CyTOF) (10) enable the characterisation of cell groups measuring the abundance for thousands of transcripts and tens of proteins at the single-cell level. The 16S rRNA and whole microbiome DNA sequencing characterise bacterial taxonomic groups (8) by probing their genetics. The relative abundance of groups of cells or microorganisms can be compared between biological or clinical conditions to identify cellular or taxonomic drivers.

Despite the importance of such comparative analyses, no method for differential variability analyses is available. Differential variability analysis is an avenue for novel discoveries through single-cell transcriptomics, such as for T-cell response in cancer (11). Also, although a wide range of differential composition methods exists, they only consider some of the five fundamental properties of cell omics-derived data (Table 1). Well known properties are (i) data is observed as counts; (ii) group proportions are compositional and negatively correlated; and (iii) the underlying proportion variability is group-specific. No available method jointly models these three properties. Log-linear proportion regression methods such as scDC (12), propeller (13) and diffcyt (14) model data compositionality (ii), and they are flexible to group-specific variability (iii) but do not model the data count distribution. Cluster-free methods such as milo (15) and DA-seq (6) are also based on log-linear models. Log-count-based methods such as MixMC (16), Bach, K. et al. (17), ANCOM-BC (18) model group-specific variability (iii) but do not model counts or data compositionality. Binomial-based methods such as Pal, Chen et al. 2021 (19) and corncob (20) model counts (i) and cell-group-specific variability (iii) but do not model the compositionality (ii). Multinomial-based methods such as ALDEx2 (21), dmbvs (22) and scCODA (23) model count data (i) and compositional properties (ii) but assume the same variability for all groups.

**Table 1.** Data properties for single-cell RNA sequencing, CyTOF and microbiome sequencing, and available methods for differential compositional analyses.

| Data properties |  |  |  |  |  |  |  |  |
| --- | --- | --- | --- | --- | --- | --- | --- | --- |
| I. | Data is observed as counts |  |  |  |  |  |  |  |
| II. | Group proportions are compositional in nature and negatively correlated |  |  |  |  |  |  |  |
| III. | The proportion variability is group-specific |  |  |  |  |  |  |  |
| IV. | Proportion mean and variability are associated |  |  |  |  |  |  |  |
| V. | Outliers are ubiquitously present |  |  |  |  |  |  |  |
| Methods | Year | Model | i | ii | iii | iv | v | Reference |
| sccomp | 2021 | Sum-constrained Beta-binomial | ● | ● | ● | ● | ● | Mangiola et al. |
| scCODA | 2021 | Dirichlet-multinomial | ● | ● |  |  |  | Buttner et al. 2021 (23) |
| quasi-binomial | 2021 | Quasi-binomial | ● |  | ● |  |  | Pal, Chen et al. 2021 (19) |
| rlm | 2021 | Robust-log-linear-model |  | ● |  |  | ● | Bach et al. (17) |
| propeller | 2021 | Logit-linear + limma |  | ● | ● | ● |  | Phipson et al. (13) |
| ANCOM-BC | 2020 | Log-linear |  | ● | ● |  |  | Lin et al. (18) |
| corncob | 2020 | Beta-binomial | ● |  | ● |  |  | Martin et al. (20) |
| scDC | 2019 | Log-linear |  | ● | ● |  |  | Cao et al. 2019 (12) |
| dmbvs | 2017 | Dirichlet-multinomial | ● | ● |  |  |  | Wadsworth et al. 2017 (22) |
| MixMC | 2016 | Zero-inflated Log-linear |  | ● | ● |  |  | Cao et al. (16) |
| ALDEx2 | 2014 | Dirichlet-multinomial | ● | ● |  |  |  | Fernandes et al. (21) |

Other important data properties have remained mostly uncharacterised. While Phipson et al. (13) introduced variability moderation through empirical Bayes (limma (24)), a mathematical description of the proportion mean-variability association (4; Table 1) across datasets and technologies is not apparent. This description would allow differential variability analysis and have profound implications for the adequacy of the very popular Dirichlet multinomial models. Similarly, the extent and the effect of outliers (v) has never been characterised for cell omic-derived data. This knowledge would guide robust method developments for cell omics data. Currently, only Bach, K. et al. (17) deal with outliers using robust regression; however, the efficacy of such an approach was not benchmarked.

Here, we introduce sccomp, a generalised method for differential composition and variability analyses based on sum-constrained independent Beta-binomial distributions. Sccomp takes into account the five main properties of cell omics compositional data. Furthermore, sccomp can simulate realistic data with the properties of any experimental dataset. The simulated data can be used to assess the adequacy of the fitted model and for benchmarking purposes. Sccomp can also transfer knowledge across datasets to improve analyses in low-data regimens.

Using sccomp on 18 datasets, we characterise the mean-variability relationship of compositional data across cell omics technologies. Our findings suggest that the Dirichlet-multinomial is inadequate for models of differential composition analysis and that incorporating the mean-variability relationship is a requirement for differential variability analysis tools. Our results also show the ubiquitous presence of outlier observations in all datasets. Using realistic simulations, we show that sccomp significantly improves performance compared to other methods. Sccomp uncovered differential microenvironmental constraints of breast cancer subtypes and cell-type-specific differences involving lymphoid and myeloid cell populations. Uniquely, the sum-constrained Beta-binomial distribution allows modelling of the compositional properties of data with mean-variability association while allowing for outlier exclusion; we anticipate its adoption in other scientific fields.

## Results

### Overview of sccomp

To take into account all five major properties of count-based compositional data (Table 1), we developed a model based on sum-constrained independent Beta-binomial distributions. Sccomp can simultaneously estimate differences in composition and variability (Figure 1C). Sccomp is compatible with complex experimental designs, including discrete and continuous covariates. The estimation is done through Hamiltonian Monte-Carlo via the Bayesian inference framework Stan (25). The hypothesis testing is performed by calculating the posterior probability of the composition and variability effects being larger than a fold-change threshold (26). The estimation is made more stable with an adaptive shrinkage in the form of a prior distribution defining the association between proportion means and variabilities (see Figure 1B and Methods). Sccomp identifies outliers probabilistically through iterative fitting (Figure 1E and Methods), which are excluded from later fits (Figure 1F).

**Figure 1.**
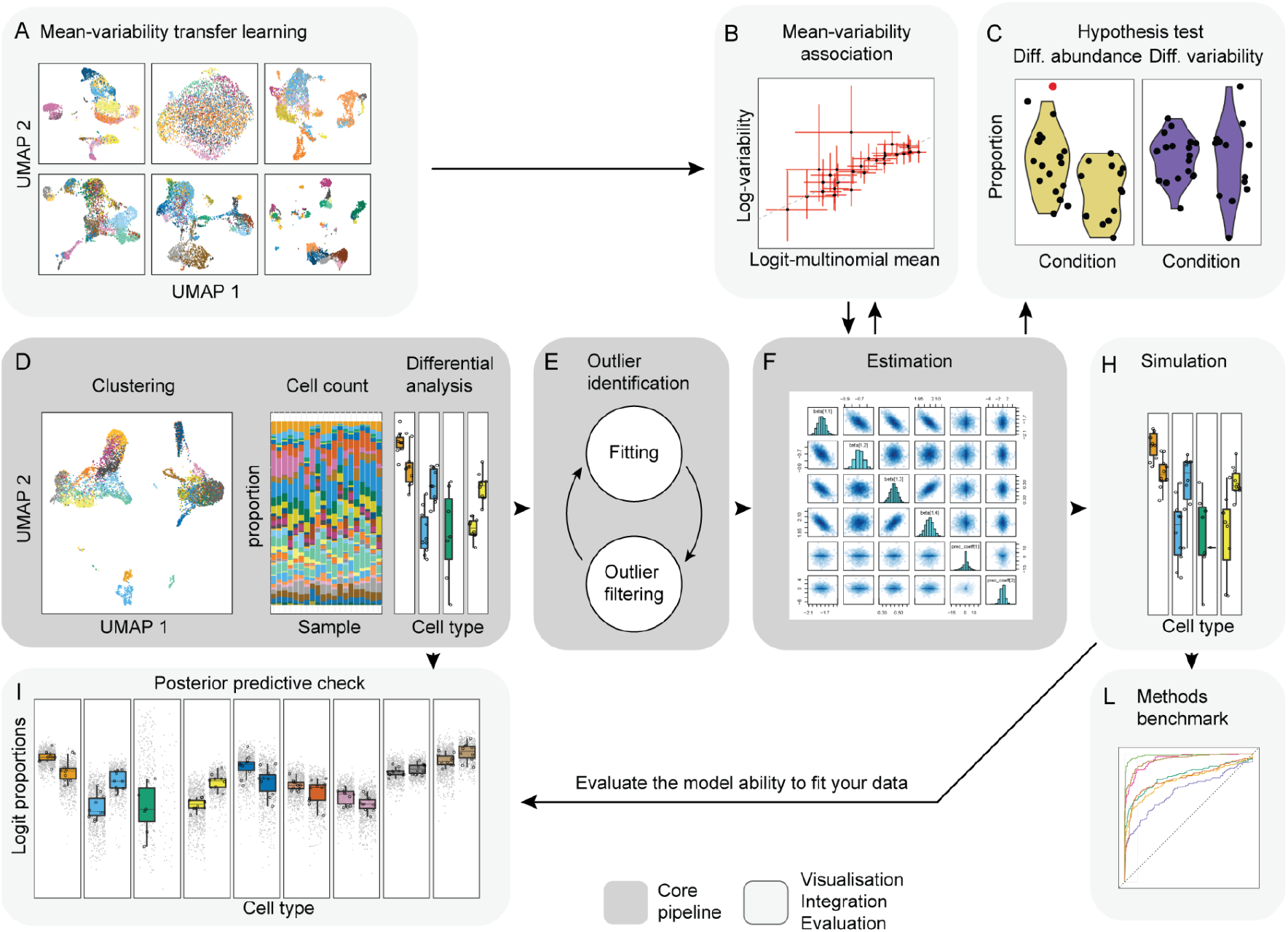
Sccomp core algorithm, data integration and visualisation. **A:** The integration of existing single-cell compositional studies gives prior information on the proportion mean-variability association (Cross-dataset learning transfer in Methods). **B:** The representation of the association between proportion means and variability (Statistical model in Methods). **C:** An example of the difference in cell-group abundance (left-hand side) and variability (right-hand side) that sccomp can estimate (Differential variability analysis in Methods). **D:** Representation of the process from cell clustering and counting that is the input for the differential composition analysis (User interface in Methods). **E:** Schematics of the iterative process of outlier identification and outlier-free data fitting that sccomp adopts for robust estimation (Iterative outlier detection in Methods). **F:** Illustration of the posterior probability distribution of regression coefficients from the model fitting (Hypothesis testing in Methods). **H:** Data simulation from the fitted model. **I:** Posterior predictive check simulates data under the fitted model and then compares these to the observed data (64) (Posterior predictive check, Methods). This check allows users to evaluate the ability of the model to fit a specific input dataset. **L:** Representation of benchmarking with realistic data that sccomp allows in a user-friendly way.

The Bayesian framework allows sccomp to incorporate the mean-variability association from other datasets (Figure 1A). This prior knowledge is helpful when only a few groups or samples are present. After fitting the model, sccomp can simulate data recapitulating the learned properties (Figure 1H). The simulated data can help identify potential failings of the model and enables benchmarking based on more realistic simulations.

### Proportion means and variabilities are log-linearly correlated in cell-omic data

To develop a method able to share information across groups, we studied the association between group proportion means and variability for 18 single-cell RNA sequencing, CyTOF and microbiome datasets (Table S1). We first fitted our sum-constrained Beta-binomial model to the data with no linear association between the logit-multinomial proportion means (μ_g_) and log-variabilities (-ω_g_) built-in (see Methods for notation), and observed the correlation of the estimates. We observed consistent positive linear homoscedastic association for all three data types (Figure 2A-left and S1, dotted line and residuals). Comparing the estimates with the mean-variability log-linear association built-in, we observe shrinkage of the variability estimates of up to 4 fold for (Figure 2A-right and S1D).

**Figure 2.**
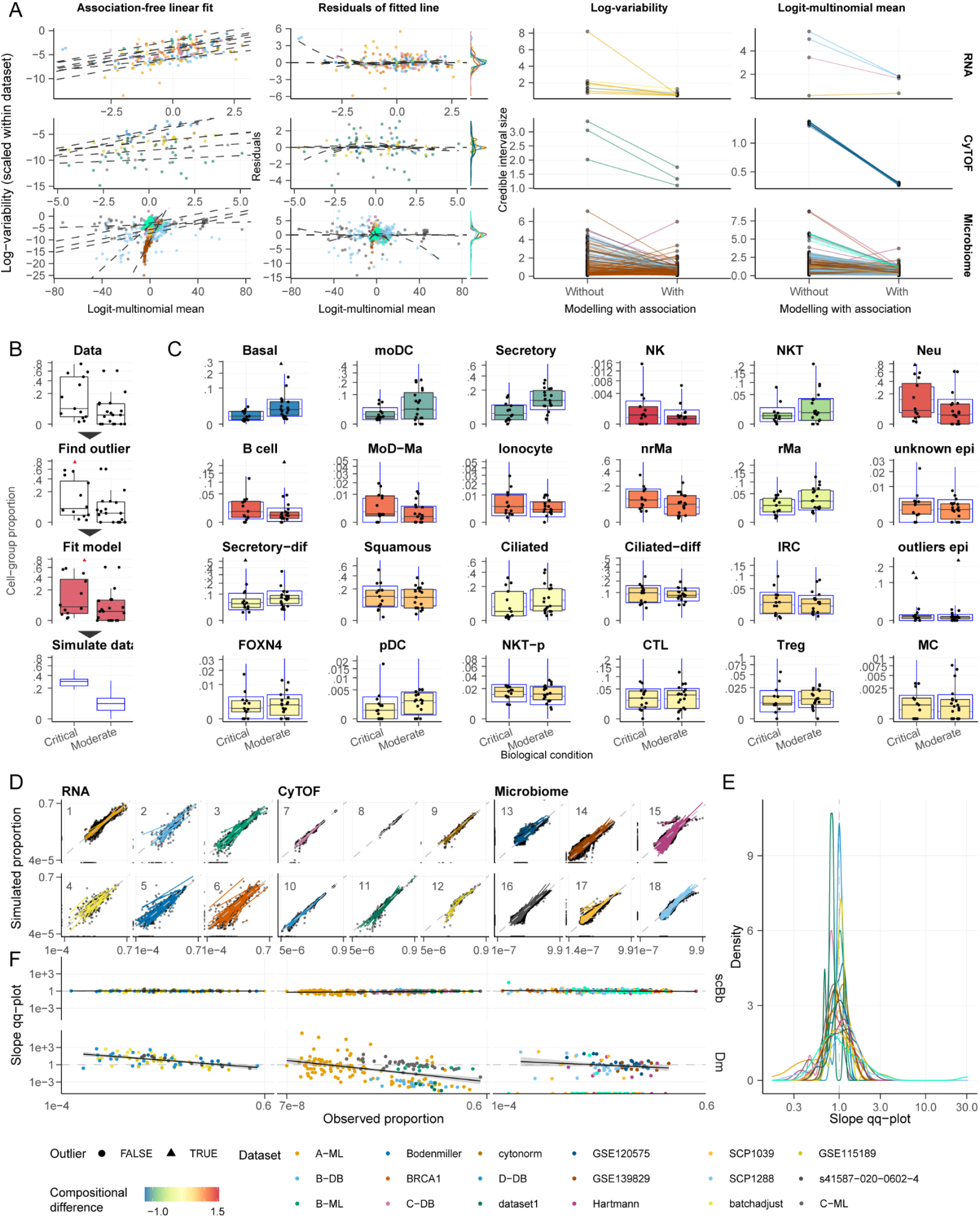
Sum-constrained Beta-binomials model mean-variability association and are adequate for experimental data from 18 studies (1–5, 17, 30–33) (Table S1), including single-cell RNA sequencing, CyTOF, and microbiome technologies. **A:** Study of the correlation between the proportion mean and variability (see Methods subsection Study of mean-variability association). The left facets refer to mean and variability estimates association with no constraints on their relationship. The dotted line is the line fitted by robust linear modelling (rlm (58)). The middle facets plot the rlm residuals versus fitted values with a lowess smoother superimposed. The facets on the right show the decrease of the size of the 95% credible intervals for all datasets. Only changes larger than 0.5 are shown (increase or decrease). **B:** The four main steps of the sccomp algorithm (see Methods section Study of model adequacy to experimental data). **C:** Example of the posterior-predictive check, with the simulated data (COVID19 dataset EGAS00001004481 (4); blue boxplots) over the observed data (colourful boxplot). The colour code expressed the magnitude of the difference estimated by sccomp across biological conditions. **D:** Scatter plot of the observed versus simulated cell-group proportions for 18 datasets, including single-cell RNA sequencing, CyTOF and microbiome (1–6). Datasets are labelled by their numeric IDs (Table S1). Each line represents a cell group. The slope of fitted lines represents the match between observed and generated data for one group (paired by their ranks), which is expected to be 1 when two distributions are the same. The dashed grey line represents a perfect linear match. **E:** The distribution of slopes of the scatter plots (panel D). **F:** Association between the slopes of the scatter plots (y-axis) and the estimated proportion abundance of each group (x-axis). Sum-constrained Beta-binomial (scBb) and Dirichlet-multinomial (Dm) are compared. If data simulated from posterior predictive distribution is similar to observed data, we expect a straight horizontal line intersecting 1.

For single-cell RNA sequencing data, modelling this association had a shrinkage effect on the variability estimates (-ω_g_; and means μ_g_ to a lesser extent), something obvious for the BRCA1 dataset for cell types with low abundance (e.g. tumour associated macrophages, Tam1, Figure S1). For CyTOF data, the shrinkage effect is evident in the Bodenmiller and cytonorm datasets. Similarly, the most significant impact can be seen for rare cell types. Microbiome data is characterised by higher uncertainty and greater spread around the regression line (before shrinkage). The impact of shrinkage there is more dramatic than in the other data types, especially for the means.

The estimated slope of the linear relationship is fairly consistent across technologies. The average slopes across datasets are 0.84, 0.47, 0.55 for single-cell RNA, CyTOF and microbiome (standard deviations 0.10, 0.22 and 0.26), respectively. Their intercepts are more variable, with the average means being −4.32, −7.19 and −5.66, and the standard deviations being 1.05, 1.86 and 5.66, respectively. Some single-cell RNA sequencing datasets show a bimodal association, where the second mode represents high-variability groups (dataset BRCA; Figure S1A). This pattern is observable in the resulting bimodal residual distribution (Figure 2A-middle and second rows of S1A, S1B and S1C). Our model uses a Gaussian mixture distribution that accurately fits both modes (Figure S1A third row; dashed lines).

### The sum-constrained Beta-binomial model achieves descriptive adequacy to experimental data across technologies

Our method can simulate realistic data based on the learned characteristics of experimental datasets (Figure 2B). This simulation is achieved by first estimating the posterior distribution from a given dataset and then generating data from the posterior predictive distribution. The posterior predictive check (27, 28) is helpful to assess the model’s descriptive adequacy (29) to specific datasets and study designs. For example, the overlay of experimental to simulated data shows the descriptive adequacy of sccomp to the COVID19 dataset EGAS00001004481 (4) (replicating quartile ranges; Figure 2C) and the absence of noticeable pathologies. To provide a more quantitative assessment, we regressed the observed and simulated data for each cell type of 18 publicly available datasets (1–5, 17, 30–33) across three cell omic technologies (Figure 2D). The fitted lines are tightly centred around the 45° reference line for all datasets (Figure 2E). This evidence suggests that proportion means and variability inference are descriptively adequate for and representative of experimental data across technologies. This trend is particularly significant considering that performing posterior predictive checks on datasets with a small sample size suffers from noise.

### The sum-constrained Beta-binomial accurately models variability across group abundance, in contrast to the Dirichlet-multinomial

Considering the existence of a mean-variability association, we assessed the ability of our model to adequately estimate the variability of small and large groups. We analysed the relationship between fitted slopes between observed and simulated proportions (Figure 2D) and the baseline abundance (estimated intercept) across 18 datasets (Table S1). We compared our model with the Dirichlet-multinomial model, a *de facto* standard for count-based compositional analyses (22, 23, 34–37).

We saw no bias in the fitted slopes of observed-simulated data across group abundance. These results present no evidence that our model underestimates or overestimates the data variability for any group, regardless of their relative abundance and the data source (Figure 2F-top). On the other hand, the Dirichlet-multinomial under/over-estimation of variability within a group is associated with group abundance (Figure 2F-bottom) for single-cell transcriptomic, CyTOF, and microbiome data. For single-cell RNA sequencing data, the variability of small groups is consistently overestimated because of the low data support (small sample size and low cell count). In contrast, for CyTOF and microbiome, where more data is available, the consistent overestimation for small groups is mirrored by an underestimation for large ones.

### The sum-constrained Beta-binomial distribution models compositionality while allowing for group-specific variability

To add dependence to a series of independent Beta-binomials, we impose a sum-to-one constraint on proportions analogously to the Dirichlet-multinomial. We hypothesised that our model would capture the compositional nature of data of a Dirichlet-multinomial, despite allowing for group-wise variability. Data was generated by a four-group Dirichlet-multinomial (with parameters 0.2, 0.6, 2.0, 4.0; Figure 3A and 3B), and the sccomp single-mean model was fitted to this data. To show the adequacy of our model of the data, we simulated data from the posterior predictive distribution. The overlay of the simulated data on the observed data shows that the densities match (red data points, Figure 3C and 5D).

**Figure 3.**
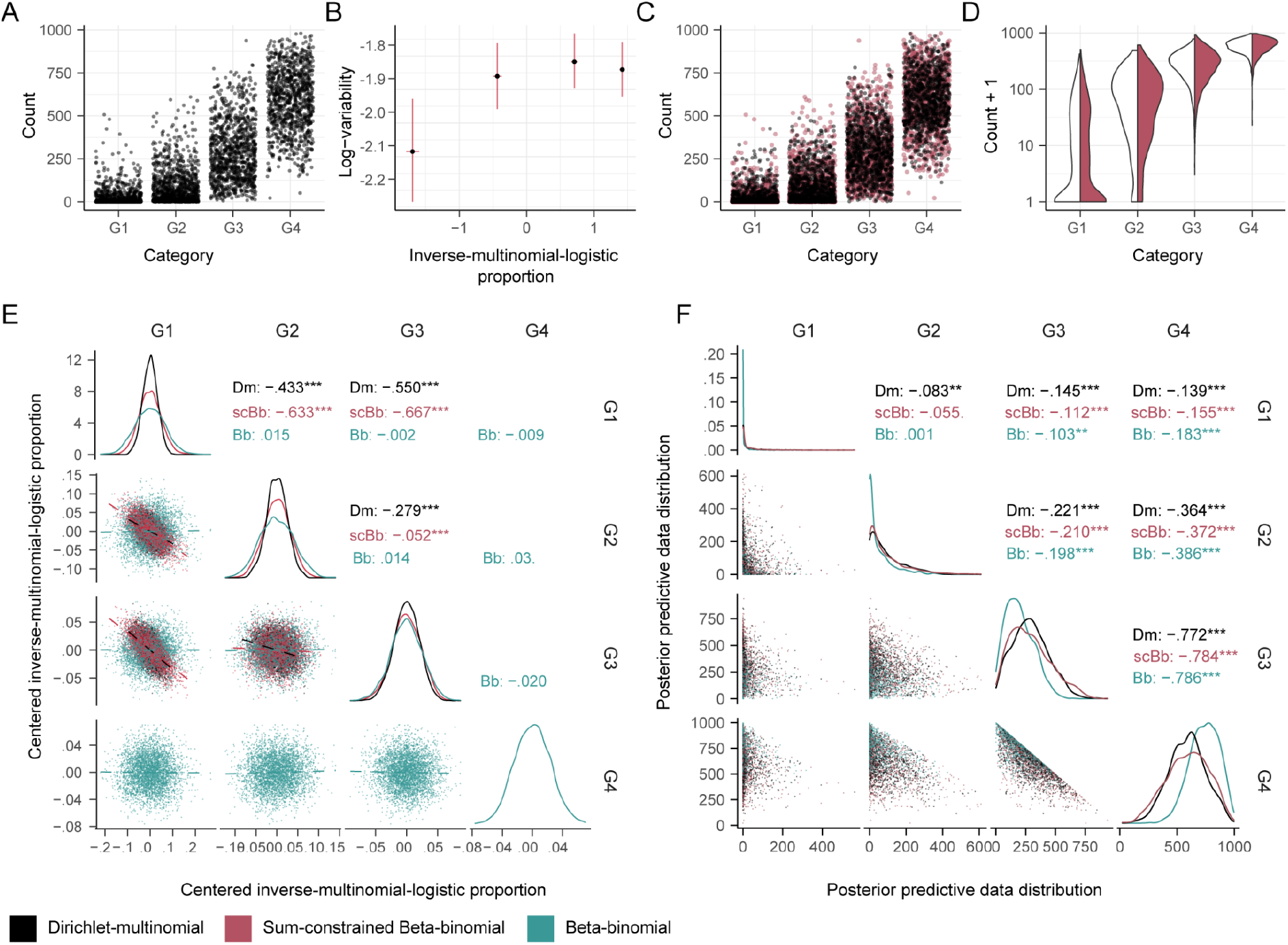
The sum-constrained Beta-binomial models the compositionality of four groups (G1, G2, G3 and G4) proportions while allowing for group-specific variability. **A:** Distribution of data simulated from a four-group Dirichlet-multinomial. **B:** Estimated mean and variability parameters from the sum-constrained Beta-binomial. The error bars represent the 95% credible intervals. **C:** Distribution of observed data (from A, black) overlaid to simulated data from the fitted model (red). **D:** Matching densities of the observed (white) and generated data (red) for the four groups. **E:** Draws from the posterior distribution of the scaled means (log-scale). The models used for estimation were Dirichlet-multinomial (Dm), (unconstrained), Beta-binomial (Bb), and sum-constrained Beta-binomial (scBb). The estimate for G4 is missing from Dirichlet-multinomial and sum-constrained Beta-binomial because it is not part of the parameter space but rather calculated as the negative sum of G1-3. The correlation is shown for each model. The stars indicate the correlation significance test (***=<0.001, calculated with GGally (65)). **F:** Overlap between the observed and generated data between each group across models.

We tested whether our model can capture the dependence structure across the proportion means, typical of compositional data, analysing the correlation among estimated means using pairs plots. We also compared the estimated means for a Dirichlet-multinomial (as a baseline) and an unconstrained (independent) Beta-binomial model. The estimated means of our model show a negative correlation structure similarly to the Dirichlet-multinomial model (Figure 3E). This correlation is strong for groups one and two (G1 and G2) and to a lesser extent for group three. On the contrary, the unconstrained Beta-binomial does not reproduce this dependence. This lack of dependence results in a higher uncertainty around the estimates, especially for low-abundance groups G1 and G2.

The differences between sum-constrained and unconstrained Beta-binomial models are reflected in the ability to simulate representative data to the Dirichlet-multinomial (Figure 3F). The marginal distributions of the predictive posterior and the weak dependence structure of the simulated data across the four groups, characteristic of the Dirichlet-multinomial, is accurately reproduced by the sum-constrained Beta-binomial. On the contrary, the unconstrained Beta-binomial generates visibly distinct data densities compared to the Dirichlet-multinomial.

### Sccomp improves the performance of differential compositional analyses

To compare the performance of sccomp with publicly available methods for differential composition analysis (Table 1), we performed a benchmarking on realistic simulated data based on the noise and outlier characteristics of the COVID19 dataset (4) (Figure 4A). The simulation was based on a logit-linear-multinomial model to ensure fairness across methods. We built a receiving-operator characteristic curve for every run and evaluated the performance using the area under the curve (AUC, up to 0.1 false-positive rates; Figure 4B).

**Figure 4.**
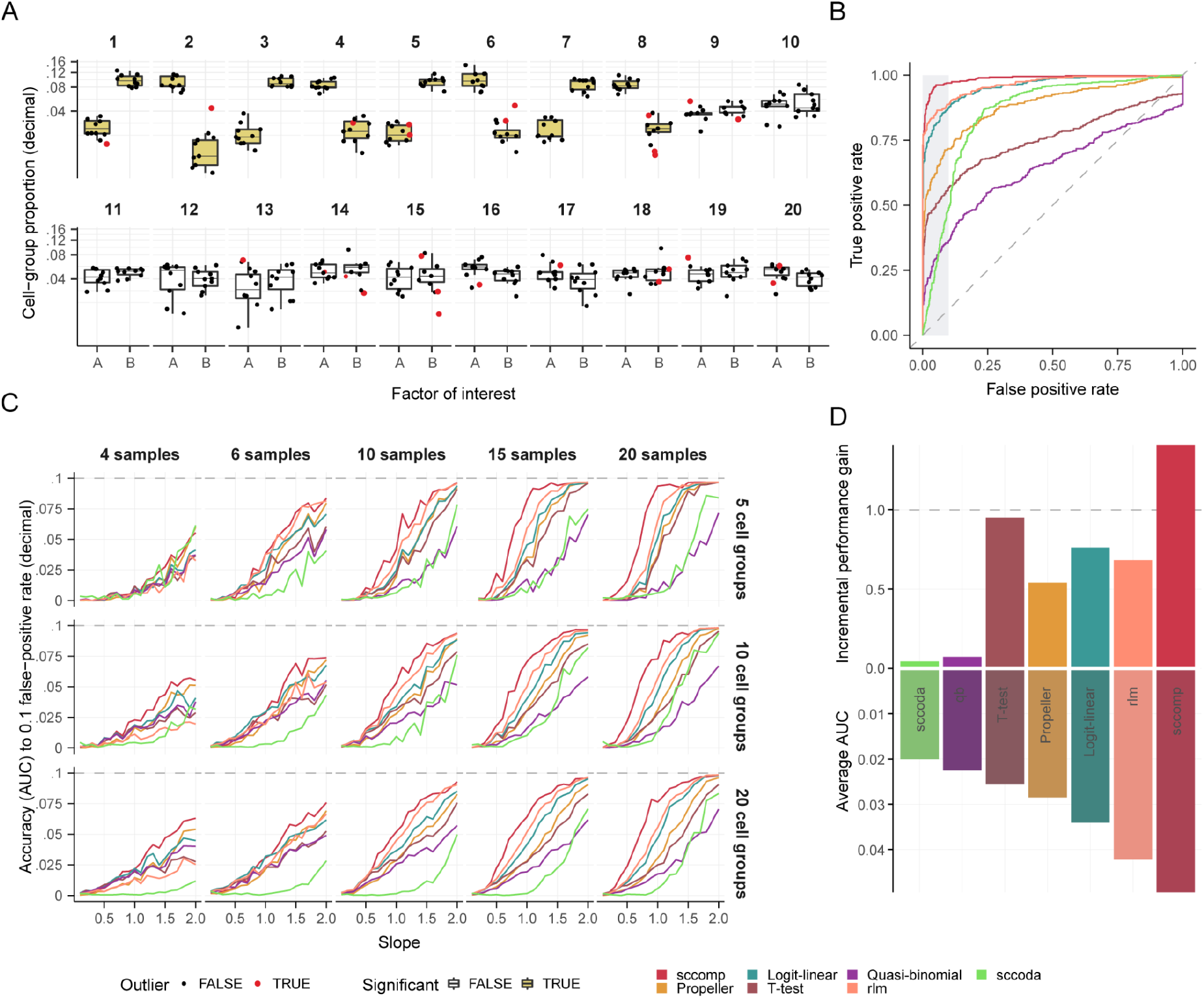
Sccomp outperforms state-of-the-art methods for realistic data simulations (including outliers) based on a logit-linear-multinomial model of the COVID19 dataset EGAS00001004481 (4) (see Methods section Benchmark). **A:** Example of a simulation with the following settings: regression slope of 1.5, 20 samples (10 per condition), 20 groups, 1000 total cells per sample, with 8 groups (40%) being differentially abundant and 12 having no differences. The yellow groups are differentially abundant. **B:** The receiving-operator characteristic (ROC) curve for the simulation in panel A, measuring the ability of the methods to identify groups as different or not based on the ground truth, as the threshold is varied. The grey area represents the false positive threshold used to calculate the area under the curve (AUC), which indicates the relative performance of each method. **C:** The comprehensive benchmarking across a range of slopes, number of samples and groups. Each performance measure represents an average of 50 areas under the curve (up to the 0.1 false-positive rate) for 50 simulations with the same parameters. **D:** Incremental performance gain across all simulation conditions (see Methods) of sccomp compared to other methods. A fold gain of 1 represents a linear incremental gain along the methods rank. Methods are ordered by their average performance across simulation conditions (bottom facet).

Overall across simulation settings, sccomp shows significantly better performance than publicly available methods, including a generic logit-linear regression (lm in R). The gain in performance tracks with the increase of slopes until the 0.1 average AUC plateau. Sccomp has the highest incremental performance gain within the method rank (Figure 4D) (1.4 folds and 1.9 fold using sum-constrained Beta-binomial-based simulations; Figure S2). It is the only method having a more-than-linear gain (i.e. > 1 fold). The method rlm and logit-linear are the second and third best performers with an incremental performance gain of 0.64 and 0.75, respectively.

Overall across simulation settings, the number of groups was the least impactful. An outlier-free benchmark (Figure S3) shows a better performance of sccomp with a smaller incremental improvement.

Sccomp can further inform the estimates by transferring information from publicly available datasets (see Cross-dataset learning transfer subsection). To test the effectiveness of this technique to regularise estimates in low-data regimens, we compared the use of uninformative or informative hyperpriors. Our results show a noticeable impact on the performance for simulations with low sample or group size, up to 2 fold (Figure S4).

### Sccomp identifies differential constraints in the microenvironments of breast cancer subtypes

We used sccomp to analyse the microenvironment of primary breast cancer from data first described by Wu et al. (3). This study analysed 26 breast cancer primary tumour tissues and identified 49 cell phenotypes. We analysed the difference in composition and variability of the triple-negative subtype (TNBC) compared to ER+ and HER2+. Our analysis led to a rich and diverse landscape of compositional and variability changes across subtypes (Figure 5A). The main feature is the depletion of cytotoxic CD8 IFN-γ, compared to HER2+ and ER+ (Figure 5B). Compared with triple-negative, the HER2+ microenvironment is enriched in several other lymphocytic populations, including CD4 follicular helper (CD4 fh in Figure 5B), CD4 CCR7+, CD4 IL7R+, T regulatory (T-reg), natural killer (NK AREG), and NKT (Figure 5B). ER+ tumours are characterised by changes in the stromal compartment, with enrichment of endothelial cells (endo ACKR1, CXCL12, RGS5) and depletion of cancer-associated fibroblasts (iCAFs2 and myCAFs4), inflammatory monocytes (Mon S100A9) and B naive cells, compared with triple-negative (Figure 5B). The differences identified by Wu et al. in the immune/stromal compartments using a t-test (3) were not labelled significant by sccomp; however, the estimated signs agree. Sccomp results are consistent with Wu et al. (3) for the enrichment of the cancer cell phenotypes (Basal, luminal, HER2+) for the respective clinical subtypes (Figure S5A).

**Figure 5.**
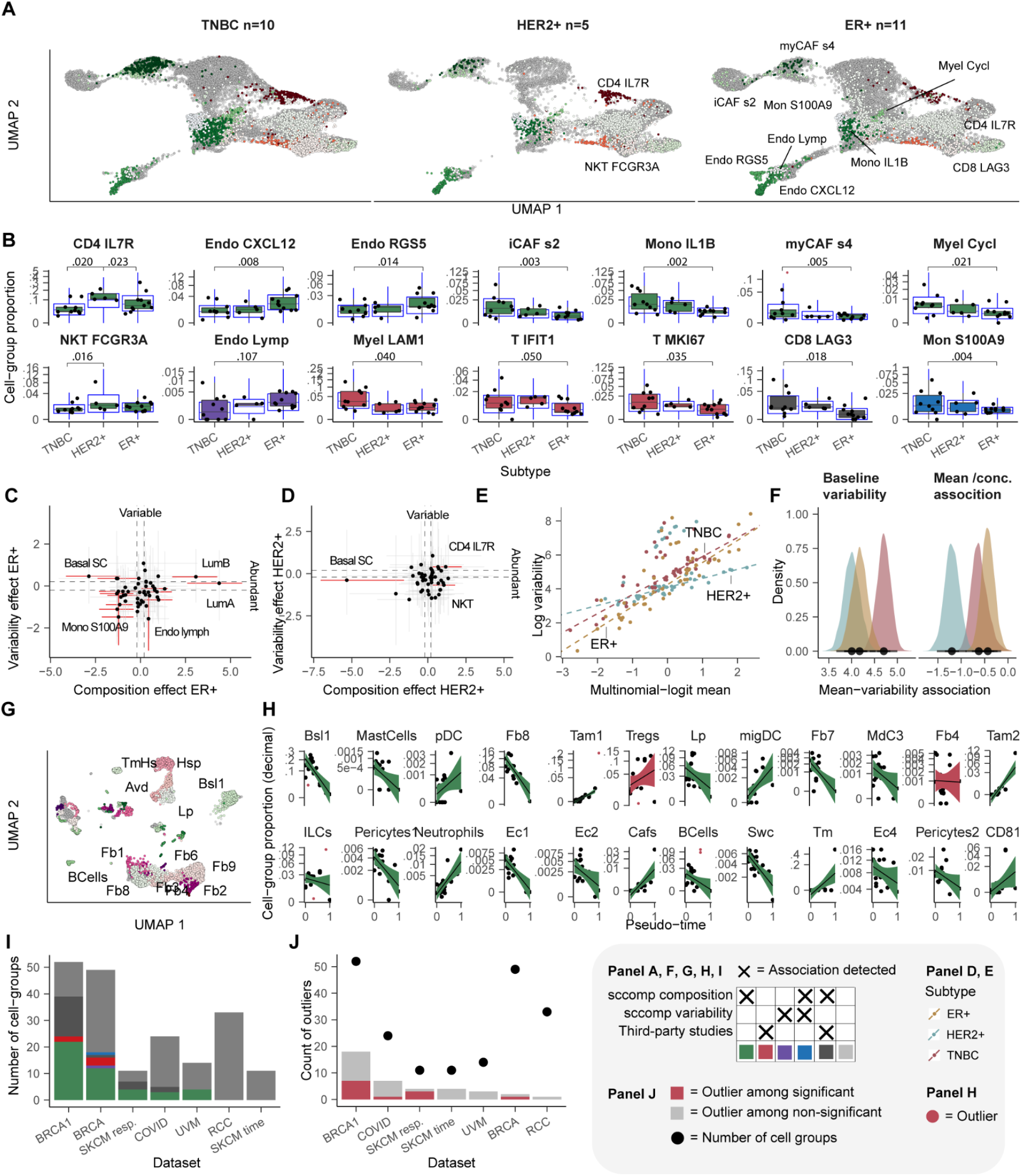
Sccomp unveiled novel results from public data from Wu et al. (3), and five single-cell RNA sequencing datasets (1, 3–5, 17). The cell groups used here were already defined in each study. **A:** UMAP projection of cells for three breast cancer subtypes. Cells are shaded according to the type of finding (e.g. green shades for novel differential composition associations). As triple-negative (TNBC) was compared with the other subtypes, only the cell groups with new findings were labelled for HER2+ and ER+ facets. **B:** Proportion distributions of the cell types with novel results (both positive and negative). The blue box plots represent the posterior predictive check. **C:** Correlation of the estimated difference in composition (x-axis) and variability (y-axis) for the triple-negative versus ER+ comparison. Error bars are the 95% credible interval. Red error bars represent significant associations. Grey dashed lines represent the minimum difference threshold of 0.2. Significant associations for cancer populations are shown in the supplementary material. **D:** Correlation of the estimated difference in composition and variability for the triple-negative versus HER2+ comparison. **E:** Mean-variability associations (in log scale) for the three cancer subtypes (see Methods subsection Reanalysis of single-cell RNA sequencing datasets). Each dot represents a cell group. The dashed lines are the sccomp estimate of such association. **F:** Posterior distributions of the intercept and slope parameters for the three subtypes, shown in panel E. **G:** UMAP projection of cells for the Bach et al. (17) dataset. Cells are shaded according to the type of finding. Only cell groups part of novel findings are labelled as text. **H:** Proportion distributions of the cell types with novel (green, red, purple, blue) and non-novel (dark and light grey) results. **I:** Count of cell groups for each dataset and the number of consistent, novel and rejected associations. The datasets are ordered by the number of novel results. **J:** Number of outliers for each dataset. Red represents outliers observation identified for differentially abundant cell groups (after outlier quarantine). Dots represent the number of cell groups per dataset. The datasets are ordered by the number of outliers identified.

Most importantly, sccomp allowed the investigation of latent microenvironmental constraints across breast cancer subtypes (see Methods subsection Reanalysis of single-cell RNA sequencing datasets). Although not immediately evident from analysing single groups (Figure 5A and 4B), the variability baseline (intercept of the mean-variability regression line; Figure 5E and 4F) for the triple-negative subtype is significantly higher than for the other cell types. This trend indicates an overall higher microenvironmental heterogeneity across patients. Also, while ER+ and triple-negative share a similar slope (−1.3 and −1.1; Figure 5A and 4B), HER2+ shows a distinct cohort-level heterogeneity profile. A markedly smaller slope indicates a more similar relative variability across cell types and potentially distinct microenvironmental processes acting for this subtype.

### Sccomp leads to novel discoveries from public datasets

To further assess the ability of sccomp in generating novel results, we expanded our analysis on a time-resolved BRCA1 model of tumorigenesis (E-MTAB-10043 (17)). Using a robust log-linear model followed by a robust F test, this study estimated 17 significant differences along the tumour developmental timeline, including fibroblast, dendritic, monocyte, and T cells. We confirmed the majority of those associations and identified 15 new associations, such as tumour-associated fibroblasts (Fb7, Fb8) and macrophages (Tam1, Tam2), neutrophils and mig dendritic cells (migDC). Five associations proposed by the study were labelled as non-significant by sccomp (Figure 5H), two of those including outliers.

To assess the usefulness of sccomp more broadly, we analysed four other single-cell RNA sequencing public datasets (Table S1). Overall, sccomp was able to generate novel results, including differential composition and variability for all datasets (Figure 5I, Figure S5B). We identified and excluded outlier observations in all datasets, with 19% of cell groups containing one or more outliers (Figure 5J). The 20% of those cell groups presented significant compositional differences after excluding outliers. The comparison between sccomp estimation and the estimation from the selected studies revealed that 15% of the calls that disagreed included one or more outliers.

## Discussion

We have introduced sccomp, a method for differential analysis of count-based compositional data. It is based on sum-constrained independent Beta-binomial distributions that share compositional characteristics with the Dirichlet-multinomial but allow group-specific variability and exclusion of outlier observation from the fit. Our model shares features with the generalised Dirichlet-multinomial (38). However, it allows for missing observations and suits outlier exclusion.

The present study describes the proportion mean-variability association for cell omic compositional data. We tested such associations across 18 single-cell RNA sequencing, CyTOF and microbiome datasets. Our results have fundamental implications. They challenge the use of the Dirichlet-multinomial distribution, a standard in count-based compositional analysis, and the use of unconstrained, independent distributions. We showed that cell omic compositional data (e.g. EGAS00001004481) with N groups can be modelled with no more than N+1 degrees of freedom (N-1 for the means and 2 for the variability). This finding implies that such unconstrained models tend to be heavily overparameterised (using 2N degrees of freedom).

Our description of mean-variability association also has implications for differential variability testing. Ignoring the mean-variability association would result in biased estimates of the differential variability necessarily associated with the differential composition estimates. Defining the correlation line in log space allowed us to disentangle differential composition and variability and provide a meaningful estimate of how cell/taxonomic proportion variability varies across samples.

While the impact of outlier observations has been approached for metagenomic data (39), our study proposes that single-cell compositional data are also outlier-rich. Our outlier identification approach overcomes the challenges of using residuals to identify outliers caused by the heteroscedastic nature of count-based compositional data and potential low sample size. We identified outliers in all single-cell RNA sequencing datasets that we have re-analysed, present in differentially and non-differentially large groups.

In real-world analyses, it is crucial to assess whether a statistical model is descriptively adequate for a specific query dataset. Sccomp offers a convenient functionality for posterior-predictive checks. Being able to generate data from a fitted model, sccomp offers a data simulation framework that reflects the properties of any target dataset while also allowing arbitrary simulation designs. Data simulation is possible using the sum-constrained Beta-binomial, Dirichlet-multinomial and logit-linear-multinomial distributions. Our realistic benchmarks (using a foreign distribution) show that sccomp confers an up to 2-fold incremental performance gain compared to previous methods.

Our reanalysis of public data demonstrates the practical application and efficacy of sccomp, which identified novel differential variability and compositional associations. We show that some of the differential composition associations proposed by the respective studies might be false-negative due to the presence of outliers. For the breast cancer dataset introduced by Wu et al. (3), we unveil differential constraints for the subtypes triple-negative, ER+ and HER2+.

This study, introducing several innovations in the field of cell omics compositional analyses such as differential variability analysis, a log-linear mean-variability relation, probabilistic outlier identification, cross-study information transfer, aims to enrich the field of single cell omics. Also, this study challenges established methodologies and provides a robust and flexible tool for the single-cell and microbiome scientific community. Being the first statistical model that fits data compositionality and group-wise variability while allowing the exclusion of outliers, we anticipate its wide adoption in other scientific fields. Sccomp is available as an R package via Bioconductor and GitHub.

## Methods

### Statistical model

The regression model underlying sccomp is based on sum-constrained independent Beta-binomial distributions. The beta distribution is a continuous probability distribution defined between zero and one. The binomial distribution is the discrete probability distribution of the number of successes obtained in a specified number of mutually independent trials, each with the same probability of success. The Beta-binomial probability distribution is the compound binomial distribution obtained when the binomial success probability is given a beta distribution. While a single Beta-binomial distribution can model the distribution of the number of elements belonging to a particular group, it cannot model the compositional nature of the data from a number of independent Beta-binomials embodied in the constraint that the underlying expected proportions belonging to the different groups must sum to one. This dependence induces a small negative correlation among the observed proportions of elements across groups, similar to that seen in the multinomial distribution. We impose this negative correlation by constraining the expected values of the group proportions (i.e. the means of our beta distributions) to sum to one.

We introduce here the common notation used in the mathematical formulation of the model. G is the number of groups, S is the number of samples, n_s_ is the total number of cells probed for sample s, k_g,s_ is the number of cells in sample s belonging to a group g. For clarity, we introduce our model in four steps. First, we describe the single-mean model; second, we describe the single-mean model with a log-linear constraint between variabilities and means; third, we introduce a two-mean model; fourth, we describe the linear model generalisation that is used in sccomp.

The Beta-binomial distribution is commonly defined using the (latent) shape parameters α and β (Equation 1) from the Beta distribution. Here and elsewhere B (α, β) denotes the classical Beta function with argument α and β. Here, we use the mean and concentration (the inverse of variability) parameterisation (π,σ) with *π*_g_ representing the mean and 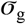 representing the concentration parameter of cell group g, this being the sum of the corresponding α and β. This parameterisation is convenient for our linear modelling. The mean is the average value of the underlying Beta distribution, while the concentration captures how concentrated the underlying Beta distribution is around its mean. The equivalence of the standard (α, β) and the alternative (π,σ) parameterisation is shown in Equations 1–3.

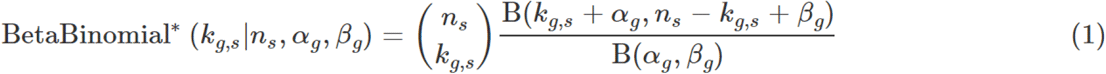

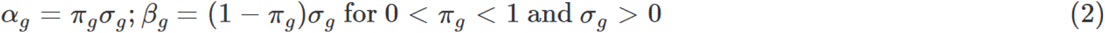

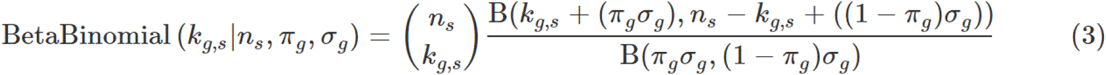

#### Step 1, Single-mean model

The parameters of the single-mean model are elements *π* = (*π*_g_) ∈ S_G+1_ (simplex) of the sum-to-one-constrained vector of size G and a vector 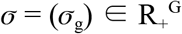 of concentrations. The data are an G×S matrix **K** = (k_g,s_) of counts, and a vector **n** = (n_s_) of length S is the sum of 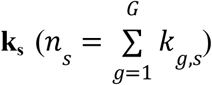. The joint probability mass function is defined by two observed quantities, **K** and **n**, depending on the parameters *π* and 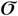, see (Equations 4–7). Statement 5 includes the sum constraint that induces the weak negative correlation of proportions characteristic of compositional data. The underlying assumption of this model is that the counts k_g,s_ from the total counts n_s_ are mutually independent Beta-binomially distributed random variables with the alternative parameters given.

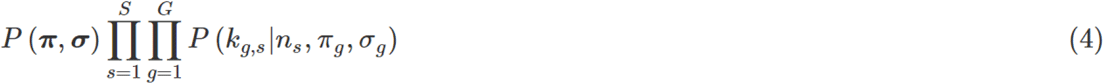

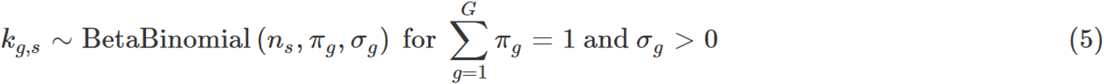

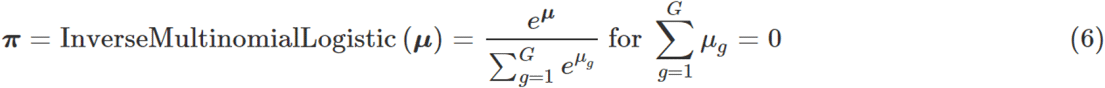

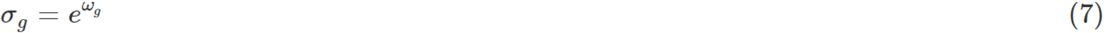

#### Step 2, Single-mean model with a (log) linear relation between concentrations and means

For this model, we transform the parameters *π* and 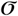 to **μ** and ω (see Equation 6 and below). The parameters *π* and 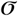 are suitable for an unconstrained single-mean model. Still, to permit a (log) linear relationship between our mean and concentration (the inverse of variability) parameters and the extension to more general linear models, we must use a different but equivalent set of parameters appropriate for linear subspaces of R^G^ The inverse-logit-multinomial (also known as *softmax*) function (Equation 6) takes a vector **μ** ∈ R^G^ and converts it into a vector of G proportions that sum to 1, the components being proportional to the exponentials of the corresponding components of **μ**. However, this mapping is many-to-one. If inverse-logit-multinomial (**μ**) = *π*, then also inverse-logit-multinomial (**μ** + c**1**_M_) = *π*, where c is any real constant and **1**_M_ is the G-vector of 1s. To make it one-to-one and so permit invertibility on its range, we need to restrict its domain. Write 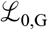 for the linear subspace of R^G^ consisting of all **μ** = (μ_g_) such that 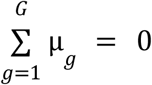. We will see that for every *π* ∈ S_G+1_ there is a unique 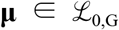 such that -I inverse-logit-multinomial (**μ**) = *π*. We call the **μ** the *logit-multinomial proportion mean* parameters, or just *mean* parameters when no confusion is likely. Letting GM denote the geometric mean, we write GM 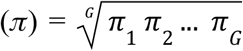. Then μ = (μ_g_) where μ_g_ = log (*π*_g_/GM (*π*)) is readily checked to satisfy our requirements, i.e. 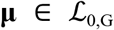 and *softmax* (**μ**) = inverse-logit-multinomial (**μ**) = ***π***. This function of ***π*** is known as its center (ed) log-ratio (CLR). We see from (7) that 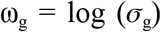, so our new parameter space is 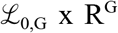. This process is also known as stick-breaking, which underlies the Dirichlet process (40, 41).

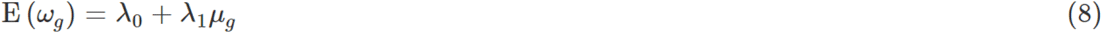

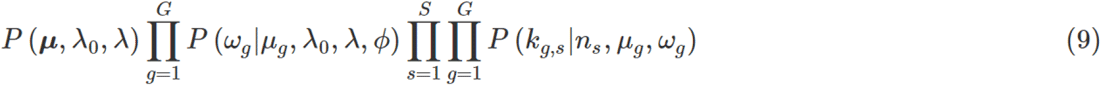

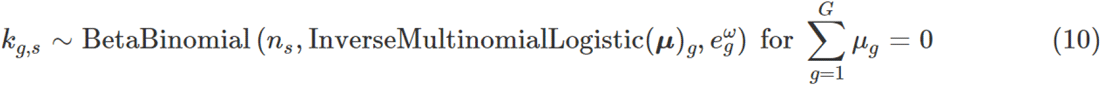

Given **μ**, the parameter ω will be given a normal prior distribution. The linear relation between **μ** and ω which underlies our development is shown in Equation (8). where λ_0_ and λ_1_ are scalars. The likelihood and priors for the single-mean model with log-linear concentration-mean relation are represented by the formulae 8–9. The complete parameter set is now 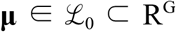, ω ∈ R^G^, λ_0_ ∈ R, λ ∈ R, and the standard deviation *ϕ* ∈ R^+^ going with the normal conditional distribution of the ω_g_ s given the μ_g_s, see (11) below. The dataset is unchanged from the original single-mean model. Before generalising this model, we introduce and use the matrix *M* = (*μ_g,s_*) of mean parameters, where *μ_g,s_* is the mean parameter for sample *s* and group *g*. The single-mean model is characterised by M having all its columns identical.

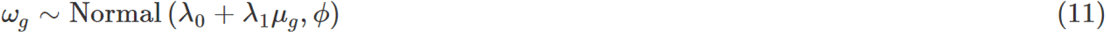

#### Step 3, Two-mean model

We now introduce the two-mean model. In this case, the matrix *M* = (*μ_g,s_*) has two potentially distinct columns, one for each of two sets of samples. For simplicity, we will call these the control and treated samples and introduce the *2*×*S* matrix X, whose 2 rows are the indicator vectors (i.e. vectors of zeros and ones) of the control and treated samples, respectively. If we now define a *G*×*2* matrix *Γ* whose columns are any two mean parameter vectors, say 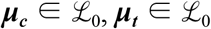, then our two-mean model has matrix *M* = *ΓX*.

#### Step 4, Arbitrary linear model

The approach of the previous paragraph can easily be generalised to arbitrary linear models. For this generalisation, we replace the *2*×*S* design matrix *X* above by an arbitrary *C*×*S* design matrix *X*, where *C* is the number of covariates associated with the samples (including one for an intercept if that is appropriate), and the *G*×*2* matrix *Γ* above becomes a general *G*×*C* matrix whose C columns are all elements of 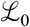. As before, *M* = *ΓX*.

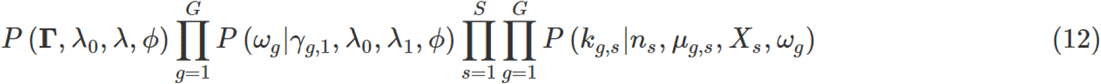

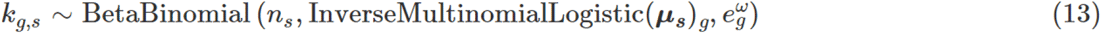

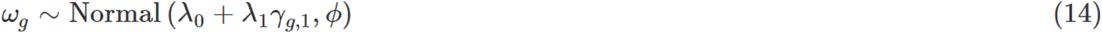

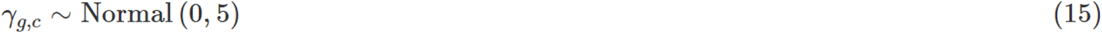

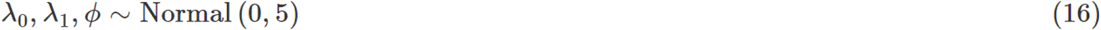

We now define the full hierarchical linear model based on the sum-constrained Beta-binomial distribution. This model is defined through the *G*×*C* parameter matrix *Γ*; ω of length *G*; *ϕ*, the scalars λ_0_ and λ_1_ and the dataset includes the *G*×*S* matrix ***K*** of counts, the *S*×*1* vector **n** of totals, and the *C*×*S* design matrix *X*. The prior normal distributions are parameterised by their means and standard deviations. *X_s_* denotes the design vector (s^th^ column) for sample s, and γ_g_ indicates the coefficient vector (g^th^ row) of *Γ* for cell-group g. Since *M* = *ΓX*, we must have *μ_g,s_* = *γ_g_X_s_*.

#### Inference

This set of sampling statements and the data (Formulae 12–16) are provided to Stan (25) to sample from a joint posterior distribution of the model parameters. Stan uses a dynamic Hamiltonian Monte Carlo sampling algorithm, a variation on the Markov-chain Monte Carlo sampling method. By default, four Markov chains are run. The number of burn-in iterations is 300 for each chain, and the number of sampling iterations is 500 per chain, giving a base of 50 draws for the 2.5% and 97.5% quantiles.

The probability of the null hypothesis (i.e. no effect across conditions) for each group is obtained by calculating the posterior probability of γ_g,c_ being larger (or smaller) than a fold-change threshold (0.2 by default). The false-discovery rate (FDR) is obtained by sorting in ascending order the probability of the null hypothesis (for any coefficient) and calculating the cumulative average as described by Stephens (26).

### Differential variability analyses

By default, the data variability is modelled with one concentration (inverse of variability) parameter ω_g_ per group (variability independent of covariates). However, the user can provide a more general variability model using a variability design matrix. For example, the concentration can be estimated conditional on a factor of interest to perform differential variability analyses. We now introduce a two-group differential variability model. The following notation is the same as in the paragraph “Step 3, Two-mean model.” of the Methods subsection “Statistical model”. As ω_g_ is the log-concentration for the cell-group g, we introduce ω_g,i_ as the concentration for cell-group g and condition i. In this model, we increase the dimensionality of ω from G to 2G, where each ω_g,1_ and ω_g,2_ represents the concentration of group g for two conditions (e.g. treatment and control). The expected value of ω for a two-group differential model and the prior distribution is described in Equations 17 and 18.

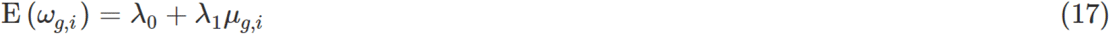

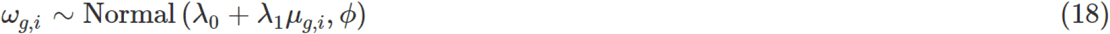

Since group proportion means and variabilities are associated (see Proportion means and variabilities are log-linearly correlated in cell-omic data) differences in composition and variability will be associated. To test the biological effects that lead to differential variability that are not explained by differences in composition, we need to subtract the contribution of differential composition from the apparent differential variability. We compute the adjusted differential variability (independent of differential composition) using Formula (19). The left side of the formula represents the (apparent) difference between variabilities, the right side of the formula represents the contribution of differential composition.

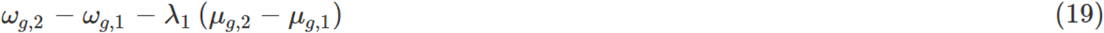

Using the dataset Wu et al., we show that without adjustment, the estimates of differential variability and composition would appear correlated (Figure S6). Often, when a cell group is differentially abundant seems also to be differentially variable. Again, this difference is the result of the mean-variability association in the first place. Without adjustment, the difference in variability would just indirectly inform us about the difference in composition without learning anything new. We show in Figure 5C and 5D that, using λ_1_ to adjust for the contribution of differential composition, we obtain estimates for differences in variability that are uncorrelated with differences in composition.

These adjusted differential variability estimates are used to carry out a test along the lines of our testing for differential composition (see Method section, Statistical model subsection).

### Iterative outlier detection

A robust iterative strategy for outlier identification was developed for negative-binomial data from bulk RNA sequencing (42). Such a strategy is necessary because a fit that includes outliers makes the model biased by definition and produces skewed estimates. Sccomp implements a three-step approach; the first two aim to identify outliers, and the third aims to estimate associations. In theory, the outlier identification process should iterate until convergence (no other outlier detected); however, our analyses across seven datasets show that two iterations always reached convergence. In the first step, the model is fitted, and posterior predictive distribution is produced for each data point from the fitted parameters. 95% credible intervals (the interval within which an unobserved parameter value falls with the probability of 0.95) are calculated from those distributions. The data points outside those intervals are labelled as outliers. This loose criterion allows (roughly because of the outlier-derived bias) 5% false-positive outliers across sample/cell-group pairs but ensures that the vast majority of outliers are identified. In the second step, the model is fitted on the outlier-free data. This fit will likely be not biased by outliers and can produce reliable posterior probability distribution to base an accurate outlier identification. The posterior predictive distribution is then produced adjusting for observation censoring (42). This adjustment is necessary because eliminating data at the distribution’s tails leads to downwards biases for the estimated variance. Credible intervals are calculated from the data distribution, allowing 5% of groups (compared to sample/cell-group pairs of the first step) to include false-positive outliers. This second step achieves a much more accurate outlier detection, for which we can better control the false-positive rate. In the third step, the model is fitted on the outlier-free datasets to estimate associations between tissue composition and biological conditions. Credible intervals of the model regression coefficients are calculated from the joint posterior distribution. For each credible interval, enough samples are drawn from the posterior distribution to provide support with 100 draws (by default). For example, for a 95% credible interval, a total of 2000 draws provides 100 draws beyond the 0.025 and 0.975 quantiles.

### Posterior predictive check

Sccomp simulates data from a specific fit to observed data using posterior predictive distribution. This data can be overlaid to the observed data to assess the model descriptive adequacy. The probabilistic framework Stan (25) is used for data simulation. The posterior distribution draws for the parameters are used to draw from one to a maximum of 2000 datasets (by default, depending on the sampling size of the fit) from the sum-constrained Beta-binomial distributions.

### Cross-dataset learning transfer

By default, our model uses uninformative gaussian hyperpriors (see the Statistical model subsection) on the intercept (λ_0_), slope (λ_1_) and standard deviation (*ϕ*) of the prior for the concentration parameter ω. **S**ccomp offers the possibility to integrate prior knowledge about the mean-variability association from other, previously analysed datasets by setting informative hyperpriors. We also provide users with a set of hyperpriors for single-cell RNA sequencing, CyTOF and microbiome data, integrating the information from the 18 analysed datasets (Table S1). We fit the model and calculate the posterior means and standard deviations of the three parameters (λ_0_, λ_1_, *ϕ*) from these data sources and set them as the mean and standard deviation of the respective hyperpriors.

### User interface

The function for linear modelling takes as input a table of cell counts (Figure 1D) with three columns containing a cell-group identifier, sample identifier, integer count and the covariates (continuous or discrete). The user can define a linear model with an input R formula, where the first covariate is the factor of interest. Alternatively, sccomp accepts single-cell data containers (Seurat (43), SingleCellExperiment (44), cell metadata or group-size). In this case, sccomp derives the count data from cell metadata. The output includes the composition and variability estimates, the probability of the effect being larger than 0.2 (by default), false discovery rate statistics, and the Markov-chain Monte Carlo convergence measures.

### Study of mean-variability association

To study the association between logit-multinomial mean *μ_g_* (where g is one cell type) and log concentration ω_g_ (negative of variability) across cell omic technologies, we gathered 7 datasets from single-cell RNA sequencing (1–5, 17, 30–33), 6 from CyTOF (45–50) and 6 from microbiome (51–56) studies (Table S1). The cell or taxonomic groups were defined in the respective studies. These datasets were analysed using the design suggested in the respective studies, assuming that the group-wise variability was independent of the covariates. For each dataset, the parameters *μ_g_* and ω_g_ were first estimated using sccomp without imposing any relationship between the two. This setting was obtained using flat, independent priors on the *μ_g_* and ω_g_. We calculated the mean, 2.5% and 97.5% quantiles from the posterior distributions of *μ_g_* and ω_g_. We then calculated the correlation between the posterior means of ω_g_ and *μ_g_* using a robust linear model (rlm, MASS (57, 58)). The residuals of the robust regression (difference between estimated ω_g_ and regression line) were calculated, and their distribution was analysed.

To assess the shrinkage effect on the concentration ω_g_ of the modelling of its linear relationship with *μ_g_*, we used sccomp including the prior of the ω_g_ given the *μ_g_*. We calculated the posterior mean and quantiles as we did with the flat independent priors. We then calculated the shrinkage as the ratio of the estimated means of *μ_g_* and ω_g_ for the two runs with or without conditional priors. We model the bimodal distribution along the regression trend present in single-cell RNA sequencing data with a mixture regression model having Gaussian distributed errors. The mixture distribution assumes an ordering of the components. The component with a higher intercept (λ_0,high_) is given a 0.9 probability, and the smaller component (λ_0,low_) is given a probability of 0.1. The slope (λ_1_) and the standard deviation (*ϕ*) are assumed to be the same for the two components (given our analyses on the single-cell RNA sequencing data with no linear association between the means and variabilities built-in). The implementation of sccomp gives the option to model the mean-variability association using mixture distribution (suggested for single-cell RNA sequencing data).

### Study of model adequacy to experimental data

To assess the adequacy (29) of the sccomp model fit to experimental data, we used the posterior predictive check (27, 28) on 7 datasets from single-cell RNA sequencing (1–5, 17, 30–33), 6 from CyTOF (45–50) and 6 from microbiome (51–56) (Table S1). For comparison, we performed the inference and analyses with both the sum-constrained Beta-binomial and the Dirichlet-multinomial models. We first used sccomp on the cell or taxonomic groups using the designs defined in the respective studies, assuming the concentrations are independent of covariates. We then used the simulation feature of sccomp to replicate those 18 datasets (i.e. posterior predictive distribution). We calculated proportion from the observed and generated counts and compared their distributions (one element being the proportion for one sample-group pair) using linear regression (lm function from R). To assess the presence of any over- or under-estimation bias conditional on the relative abundance, we compared the slope of the association between observed and generated data with the baseline group abundance (intercept coefficient).

### Reanalysis of single-cell RNA sequencing datasets

To assess the ability of sccomp to generate discoveries from publicly available datasets, we applied sccomp to 6 single-cell RNA sequencing datasets (1–5, 17). The cell groups were defined in the respective studies, and the concentration was set to be conditional to the factor of interest (first covariate in the formulae in Table S1). We then compared the significant associations and the presence of outliers identified by sccomp with the findings presented in the respective studies. We estimated the differential composition for all datasets, while only for the datasets with a binary factor of interest and a sample size larger than 10, we estimated the differential variability.

For the BRCA dataset by Wu et al. (3), we went beyond the cell group composition/variability analysis to a higher-level inference. We can think about the mean-variability association as a signature specific to a group of samples (e.g. cancer tissue from patients with a triple-negative breast cancer subtype). The intercept of the regression line represents the baseline variability within a tissue across samples. The slope of the regression line represents the uniformity of the variability across cell types. A smaller slope indicates that all groups (cell-types) are similarly variable; a larger slope indicates that the larger groups are relatively much more variable than small groups. We can compare these high-level properties across groups of samples, for example, from different breast cancer subtypes. This analysis can identify different high-level signatures that might underlie different biological constraints and processes. For that, we ran sccomp independently for the samples according to the factor of interest “subtype” and compared the posterior distribution of the estimated mean-variability association (λ_1_) and the baseline variability (modelled in its negative form as concentration λ_0_).

### Benchmark

We base our benchmark on simulated data with realistic characteristics based on the COVID19 dataset EGAS00001004481 (4). Data was simulated based on a logit-linear-multinomial model to ensure fairness across methods. For the simulation, we calculated group proportions across samples and fitted a logit-linear regression model using Stan (25). This model captures the mean-variability association similar to the default sccomp model. We use the posterior distribution to simulate the benchmark datasets, replacing the intercepts and slopes with given ones to establish ground truth. For each simulation, we randomly selected 40% of the groups to be compositionally different between conditions. With the values of the specified parameters, we simulated data from a logit-linear-multinomial model to obtain counts. The total cell count for each sample was set to 1000.

We based the simulation on three variables: the magnitude of the difference, number of samples per condition, and number of groups. The outliers were injected with realistic frequency (10%) and magnitude (from 2 to 10 fold increase or decrease), observed in our data-integration analyses. For each simulation, we produced a receiving-operator characteristic (ROC) curve ranking groups by their statistics and comparing them with the ground truth of significant/non-significant differentially abundant groups. We calculated the area under the curve (AUC) from the receiving-operator characteristic curves to a 0.1 false-positive rate (grey shade in Figure 4B). For each combination of the simulation parameters, we simulated 50 datasets and averaged across the areas under the curves. For comparative purposes, we performed a benchmark (as described above) simulating data without outliers and using the sum-constrained Beta-binomial distribution.

We assessed the leap of performance improvement that sccomp provides compared to its next-best in relation to the performance gain of all methods. We call this measure incremental performance gain. An incremental performance gain of one indicates that, compared to the average performance improvement that any method had with its second-best, a method had the same improvement. This can be thought of as linear incremental performance gain. For each of the 15 simulation settings, we calculated the average area under the ROC curve (AUC) to obtain a unique performance score. From this average, we excluded the slope ranges where the performance of most algorithms approached a plateau to calculate the difference in performance in the most informative simulation regimens. For each simulation setting, we ranked methods based on the performance score. We calculate the gain in performance as the difference in average AUC between each method and their next-best ranked (e.g. first against second, second against third). The fold gain in performance was calculated as the ratio between the gain in performance of each method and the average of all others.

### Data analysis and manipulation

The data analysis was performed in R (59). Data wrangling was done through tidyverse (60). Single-cell data analysis and manipulation were done through Seurat (43) (version 4.0.1), tidyseurat (61) (version 0.3.0), and tidybulk (62) (version 1.6.1). Parallelisation was achieved through makeflow (63). Pair plots created with GGally (cran.r-project.org/web/packages/GGally).

## Code availability

Sccomp is implemented as an open-source package in R (Github: stemangiola/sccomp) and is installable from Bioconductor. Code used to generate figures and perform analyses can be found at https://github.com/stemangiola/sccomp/tree/master/dev.

## Acknowledgements

We thank Davis MacCarthy, Gordon Smyth and Jeffrey Pullin for the fruitful discussions about the method. We thank all the Stan community for the constant support.

## Ethics declarations

Authors have no competing interests to declare.

## Contributions

S.M. conceived, designed and implemented the methods and performed the analyses under the equal supervision of A.T.P., T.P.S. and H.S. A.S., M.T. and E.Z. contributed equally to the analyses of CyTOF and microbiome data. M.M. and Z.G. contributed to the analyses of single-cell RNA sequencing data. A.F.R. contributed to the benchmark. All authors contributed to the writing of the article.

## Funding

S.M. was supported by the Victorian Cancer Agency Early Career Research Fellowship (ECRF21036). A.T.P. was supported by an Australian National Health and Medical Research Council (NHMRC) Senior Research Fellowship (1116955). S.M. and A.T.P. were supported by the Lorenzo and Pamela Galli Next Generation Cancer Discoveries Initiative. The research benefited from the Victorian State Government Operational Infrastructure Support and Australian Government NHMRC Independent Research Institute Infrastructure Support.

## Supporting information

**Table S1.**
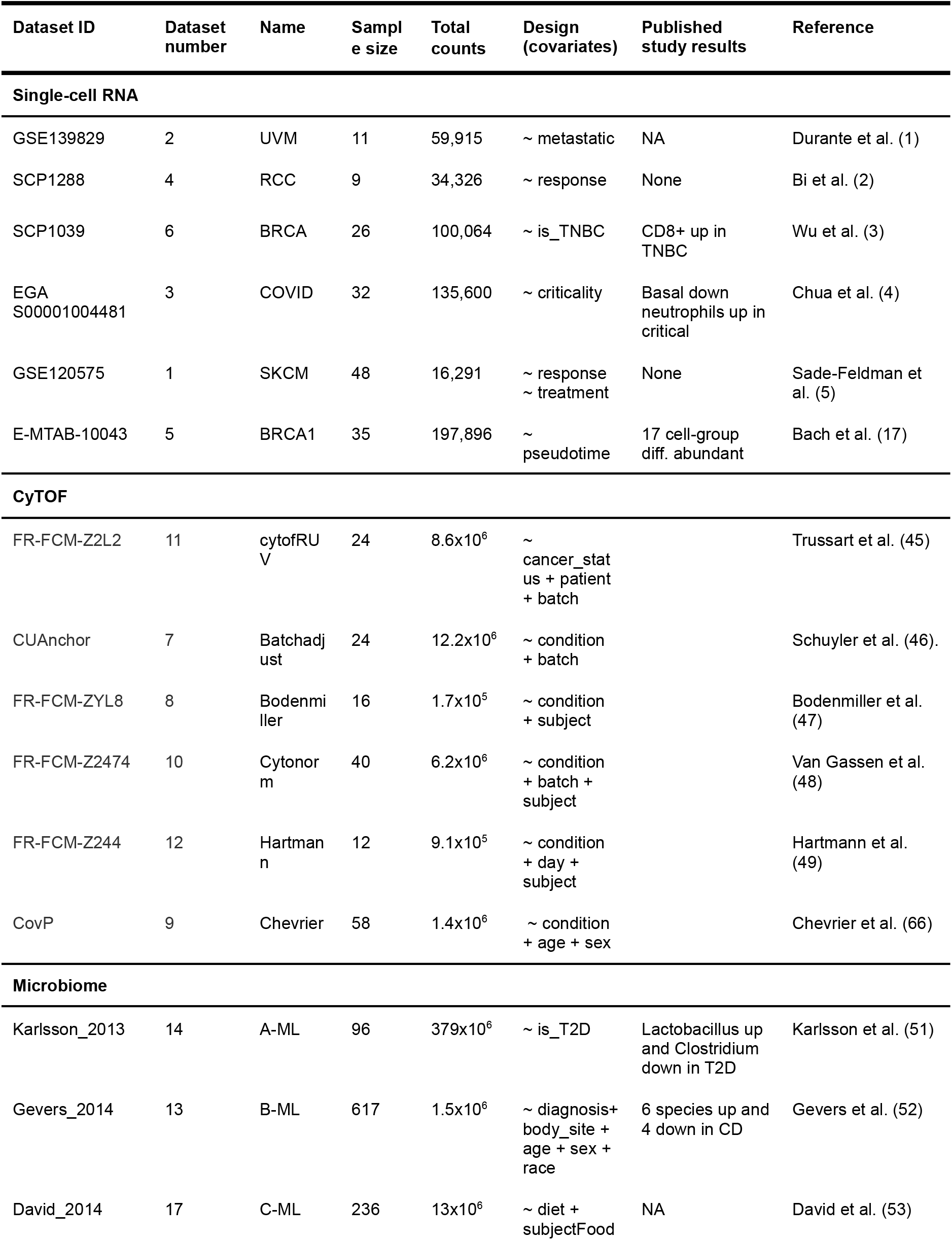
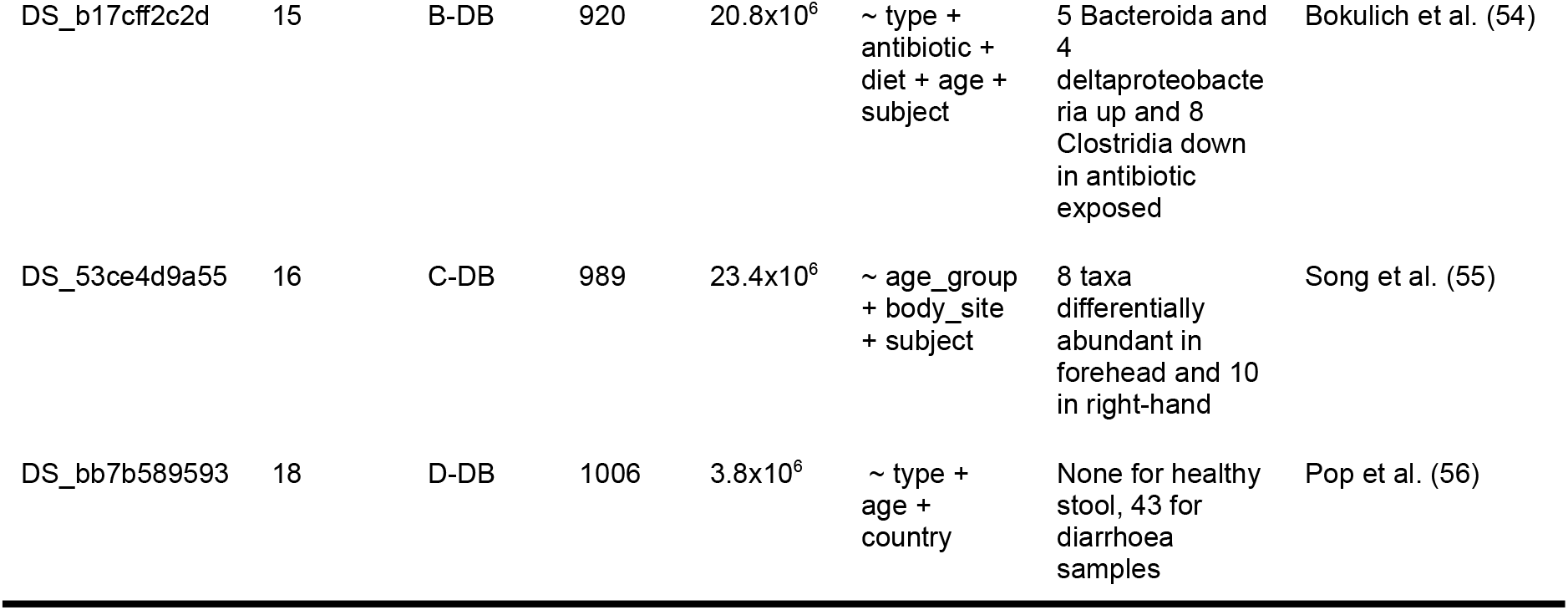
Publicly available datasets and studies used for re-analyses. Name refers to the abbreviations used in the article. Multiple analysis designs tested for a study are represented by multiple rows in the Design column. NA in the Design or Published study results is used if the published study performed no differential composition. None refer to no significant association detected by a study.

**Figure S1.**
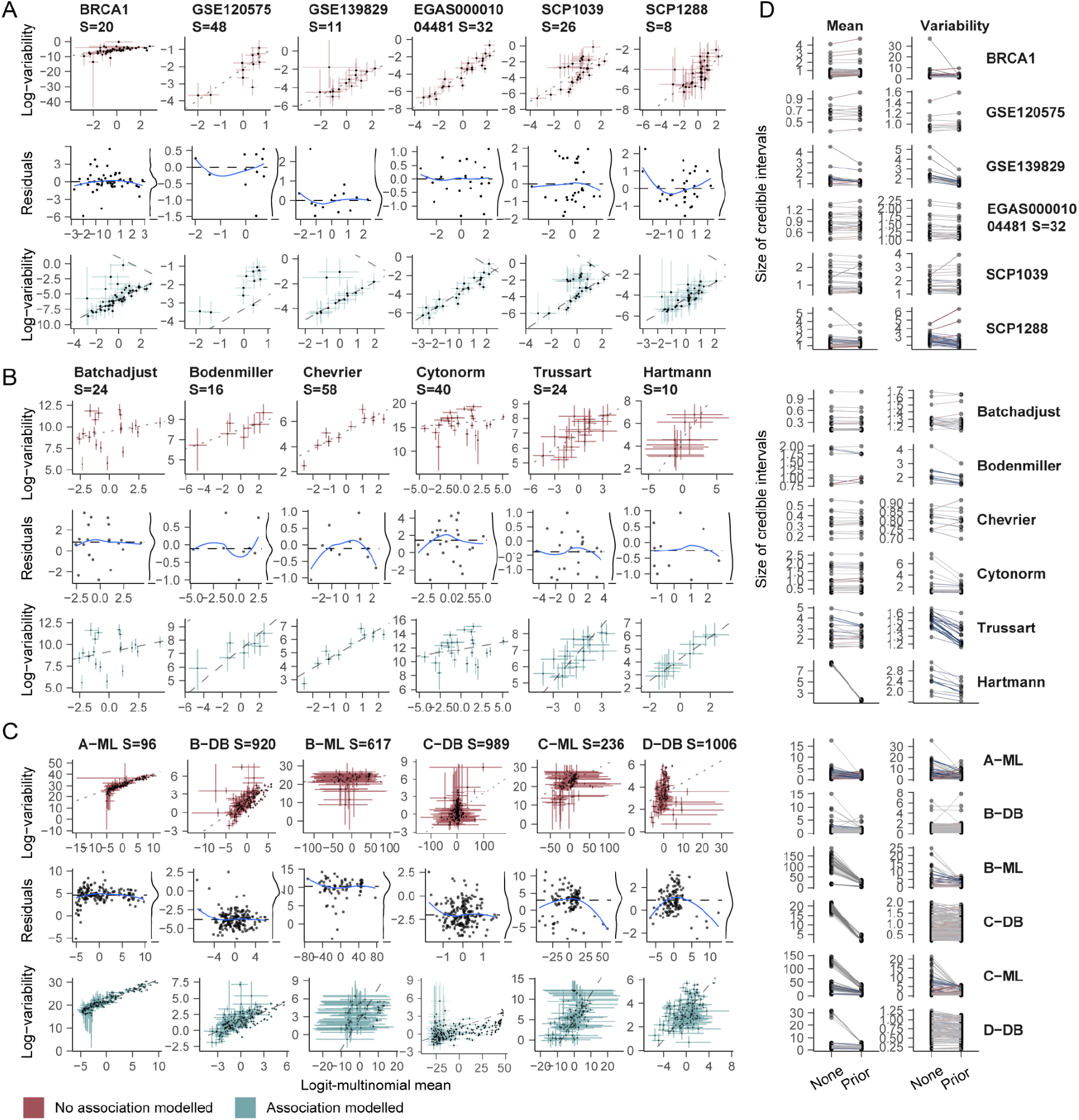
Study of the correlation between the proportion mean and variability from 18 datasets (1–5, 17, 30–33) (Table S1, see Methods subsection Study of mean-variability association). For panels A, B and C, the points are the posterior means of the parameters. The error bars are the 95% credible intervals. The first row refers to mean and variability estimates association with no constraints on their relationship. The dotted line is the line fitted by robust linear modelling (rlm (58)). The second row has rlm residuals vs fitted values with a (blue) lowess smoother superimposed. The third line represents the estimates with the mean-variability association modelled. The dashed lines are the correlation estimated by sccomp. **A:** Mean-variability association for the single-cell RNA sequencing data. For this data type, the bimodal association is modelled (see Methods, Study of mean-variability association) **B:** CyTOF data. The association is modelled as unimodal. **C:** Metagenomics data. The association is modelled as unimodal. **D:** The change in the size of the 95% credible intervals without/with constraints on the mean-variability relationship (None/Prior) for all datasets.

**Figure S2.**
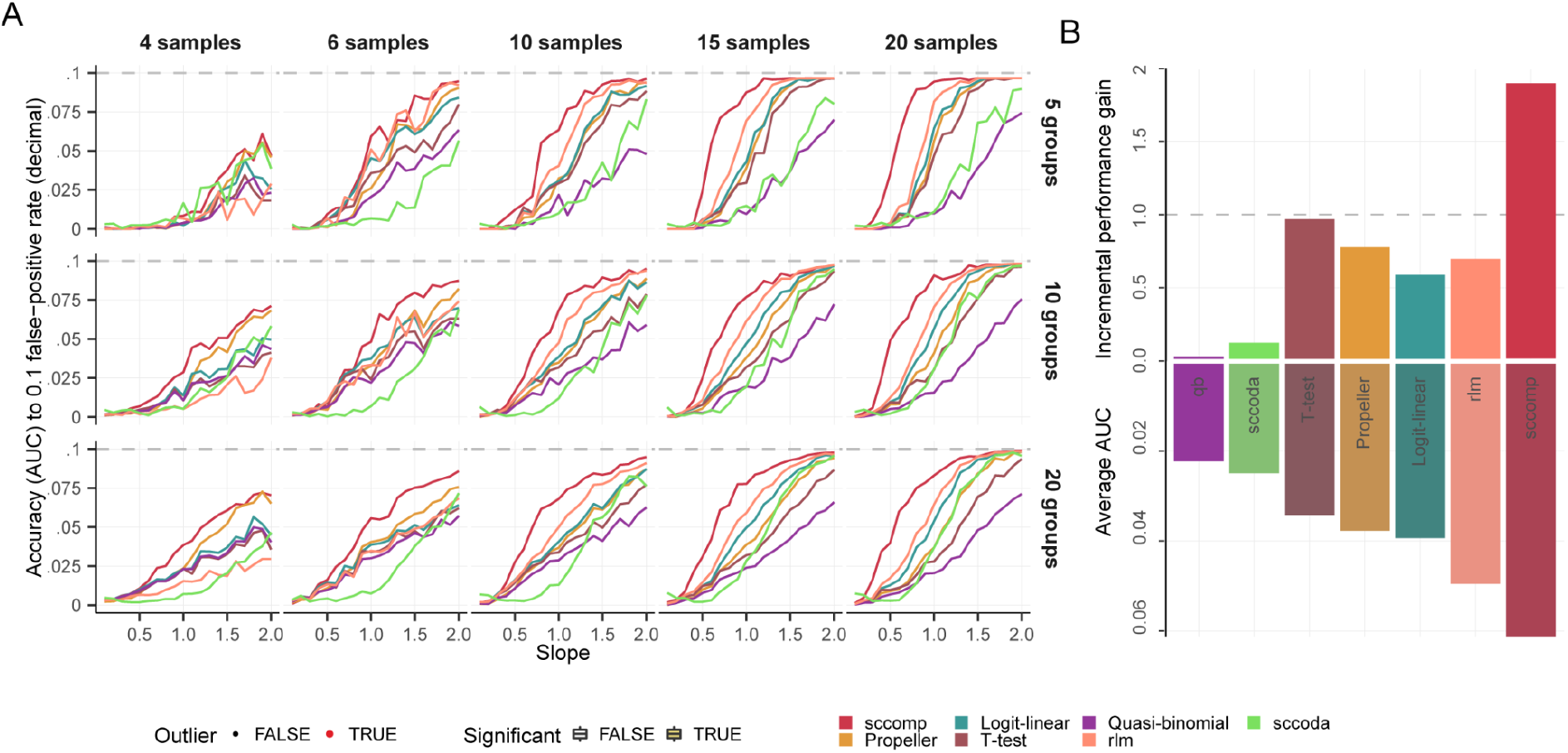
Benchmark on realistic simulated data from the COVID19 dataset EGAS00001004481 (4) using the sum-constrained Beta-binomial model.

**Figure S3.**
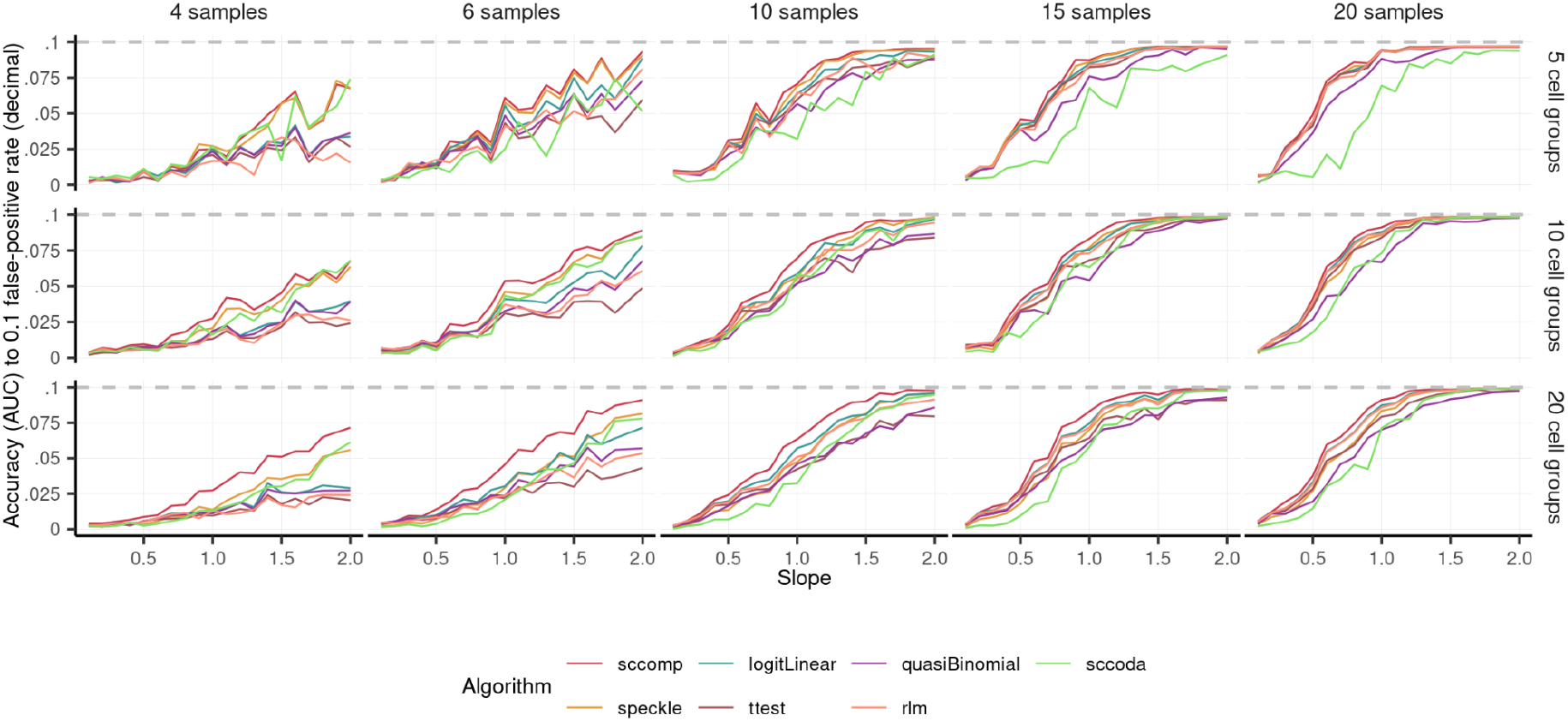
Benchmark on realistic simulated data from the COVID19 dataset EGAS00001004481 (4) using the outlier-free logit-linear-multinomial model. The comprehensive benchmark across a range of slopes, number of samples and groups. Each performance measure represents an average of 50 areas under the curve (up to the 0.1 false-positive rate) for 50 simulations with the same parameters.

**Figure S4.**
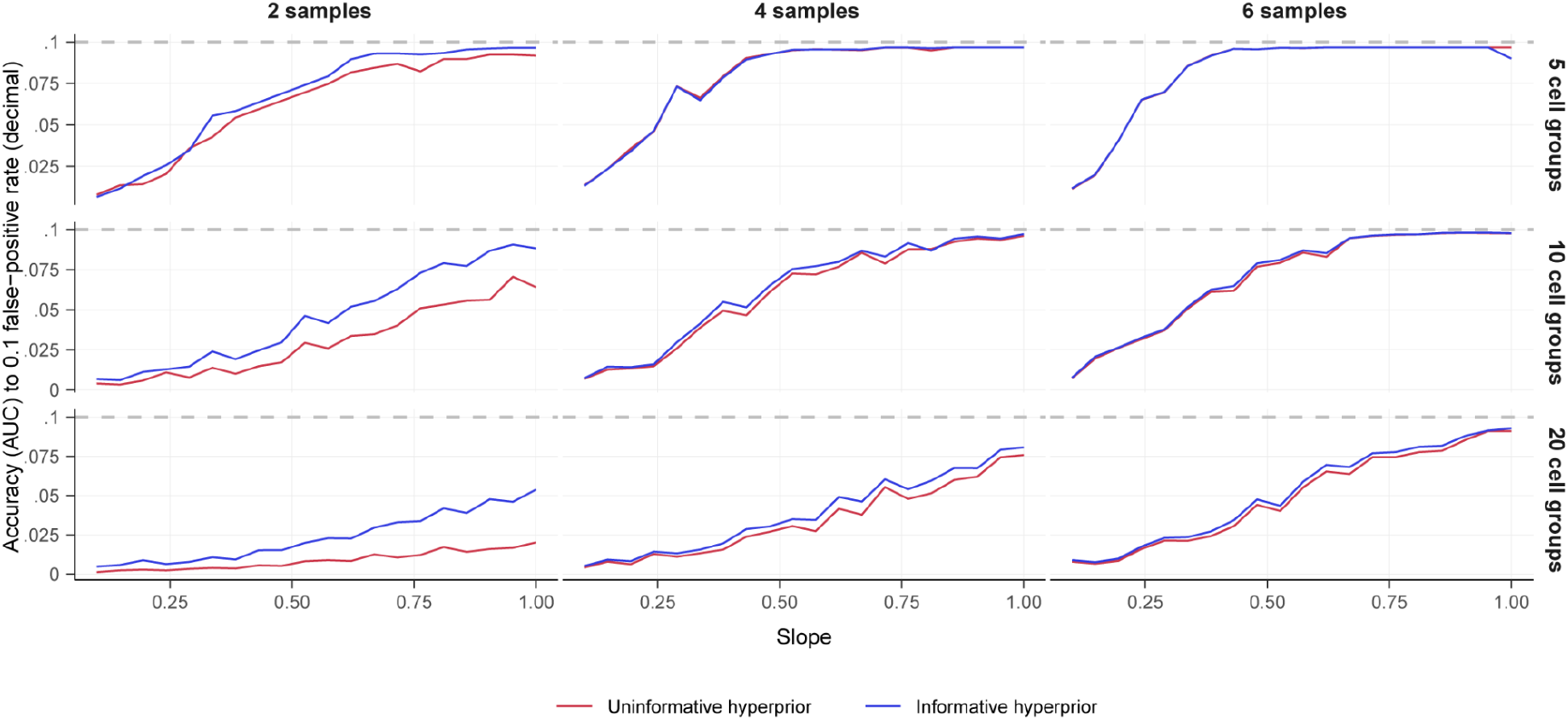
Benchmark for sccomp comparing the use of cross-dataset learning transfer in a low-sample and low-group size regimen.

**Figure S5.**
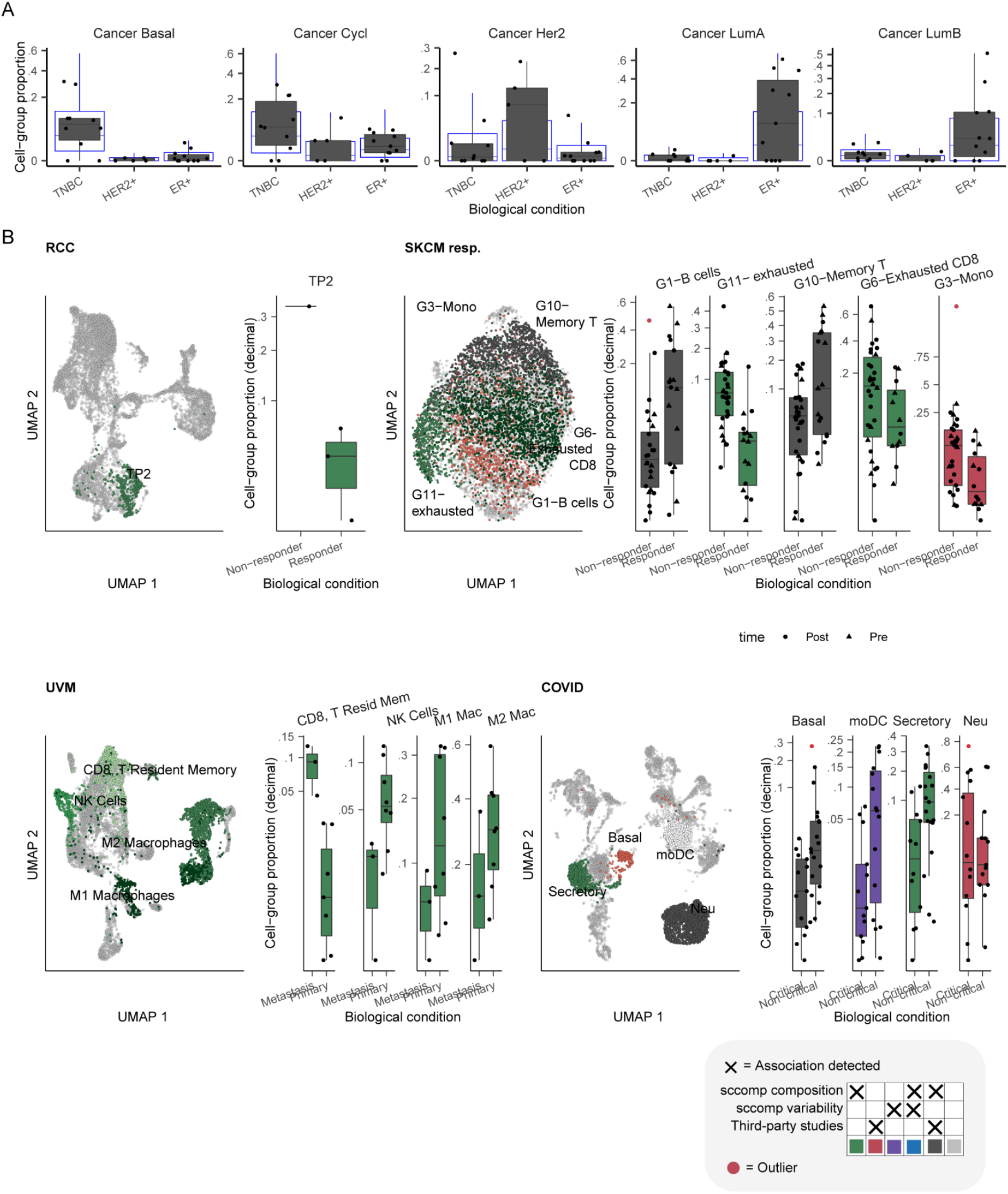
Application of sccomp of publicly available dataset. This figure is a companion to Figure 5. **A:** Proportion distributions of the cell types with the detected associations for cancer populations for the dataset Wu et al. (3). The blue box plots represent the posterior predictive check. **B:** UMAP projection of cells for three breast cancer subtypes and boxplots for other datasets (Table S1). Cells are shaded according to the type of finding (e.g. green shades for novel differential composition associations).

**Figure S6.**
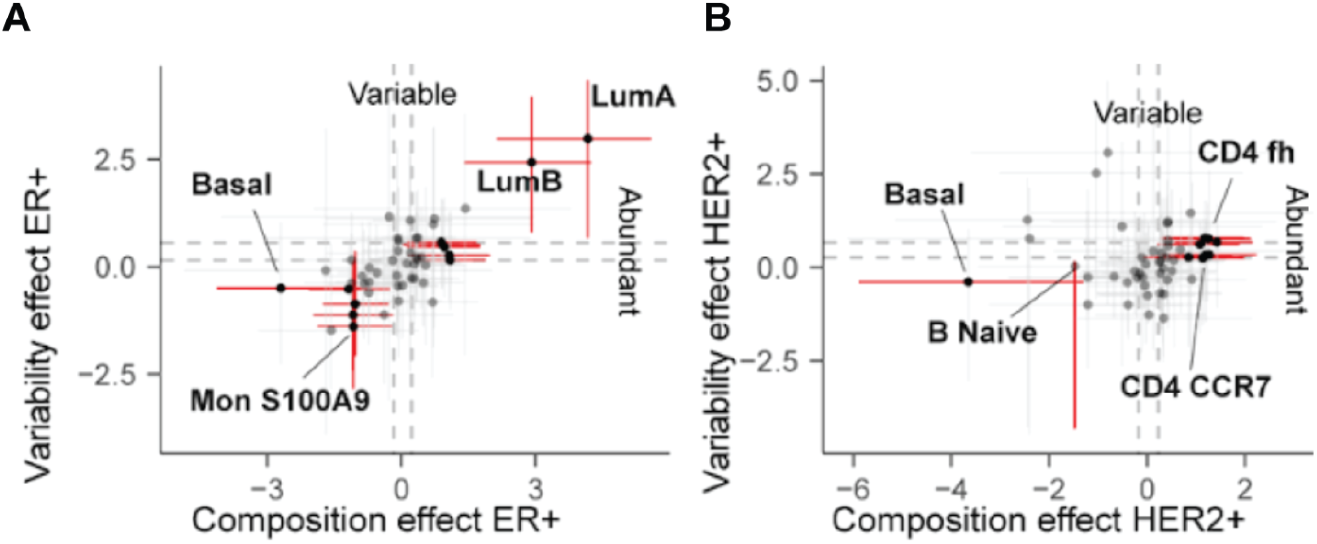
A counterpart of Figure 5C and 5D (see Methods subsection Differential variability analyses), without adjustment for the mean-variability association. **A:** Estimated difference in composition (x-axis) and variability (y-axis) for the triple-negative versus ER+ comparison, without adjusting the mean-variability association. Error bars are the 95% credible interval. Red error bars represent significant associations. Grey dashed lines represent the minimum difference threshold of 0.2. Significant associations for cancer populations are shown in the supplementary material. **B:** Estimated difference in composition and variability for the triple-negative versus HER2+ comparison without adjusting the mean-variability association.

## Notes

https://github.com/stemangiola/sccomp

### Competing Interest Statement

The authors have declared no competing interest.

## References

1. M. A. Durante, et al., Single-cell analysis reveals new evolutionary complexity in uveal melanoma. Nat. Commun. 11, 496 (2020).

2. K. Bi, et al., Tumor and immune reprogramming during immunotherapy in advanced renal cell carcinoma. Cancer Cell 39, 649–661.e5 (2021).

3. S. Z. Wu, et al., A single-cell and spatially resolved atlas of human breast cancers. Nat. Genet. 53, 1334–1347 (2021).

4. R. L. Chua, et al., COVID-19 severity correlates with airway epithelium-immune cell interactions identified by single-cell analysis. Nat. Biotechnol. 38, 970–979 (2020).

5. M. Sade-Feldman, et al., Defining T Cell States Associated with Response to Checkpoint Immunotherapy in Melanoma. Cell 175, 998–1013.e20 (2018).

6. J. Zhao, et al., Detection of differentially abundant cell subpopulations in scRNA-seq data. Proc. Natl. Acad. Sci. U. S. A. 118 (2021).

7. Y. Fan, O. Pedersen, Gut microbiota in human metabolic health and disease. Nat. Rev. Microbiol. 19, 55–71 (2021).

8. A. L. Byrd, Y. Belkaid, J. A. Segre, The human skin microbiome. Nat. Rev. Microbiol. 16, 143–155 (2018).

9. M. Karlsson, et al., A single-cell type transcriptomics map of human tissues. Sci Adv 7 (2021).

10. R. K. Cheung, P. J. Utz, Screening: CyTOF-the next generation of cell detection. Nat. Rev. Rheumatol. 7, 502–503 (2011).

11. E. F. Davis-Marcisak, et al., Differential Variation Analysis Enables Detection of Tumor Heterogeneity Using Single-Cell RNA-Sequencing Data. Cancer Res. 79, 5102–5112 (2019).

12. Y. Cao, et al., scDC: single cell differential composition analysis. BMC Bioinformatics 20, 721 (2019).

13. B. Phipson, et al., propeller: testing for differences in cell type proportions in single cell data. bioRxiv, 2021.11.28.470236 (2021).

14. L. M. Weber, M. Nowicka, C. Soneson, M. D. Robinson, diffcyt: Differential discovery in high-dimensional cytometry via high-resolution clustering. Commun Biol 2, 183 (2019).

15. E. Dann, N. C. Henderson, S. A. Teichmann, M. D. Morgan, J. C. Marioni, Differential abundance testing on single-cell data using k-nearest neighbor graphs. Nat. Biotechnol. (2021) https://doi.org/10.1038/s41587-021-01033-z.

16. K.-A. Lê Cao, et al., MixMC: A Multivariate Statistical Framework to Gain Insight into Microbial Communities. PLoS One 11, e0160169 (2016).

17. K. Bach, et al., Time-resolved single-cell analysis of Brca1 associated mammary tumourigenesis reveals aberrant differentiation of luminal progenitors. Nat. Commun. 12, 1502 (2021).

18. H. Lin, S. D. Peddada, Analysis of compositions of microbiomes with bias correction. Nat. Commun. 11, 3514 (2020).

19. B. Pal, et al., A single-cell RNA expression atlas of normal, preneoplastic and tumorigenic states in the human breast. EMBO J. 40, e107333 (2021).

20. B. D. Martin, D. Witten, A. D. Willis, MODELING MICROBIAL ABUNDANCES AND DYSBIOSIS WITH BETA-BINOMIAL REGRESSION. Ann. Appl. Stat. 14, 94–115 (2020).

21. A. D. Fernandes, et al., Unifying the analysis of high-throughput sequencing datasets: characterizing RNA-seq, 16S rRNA gene sequencing and selective growth experiments by compositional data analysis. Microbiome 2, 15 (2014).

22. W. D. Wadsworth, et al., An integrative Bayesian Dirichlet-multinomial regression model for the analysis of taxonomic abundances in microbiome data. BMC Bioinformatics 18, 94 (2017).

23. M. Büttner, J. Ostner, C. L. Müller, F. J. Theis, B. Schubert, scCODA is a Bayesian model for compositional single-cell data analysis. Nat. Commun. 12, 6876 (2021).

24. G. K. Smyth, “limma: Linear Models for Microarray Data” in Bioinformatics and Computational Biology Solutions Using R and Bioconductor, R. Gentleman, V. J. Carey, W. Huber, R. A. Irizarry, S. Dudoit, Eds. (Springer New York, 2005), pp. 397–420.

25. B. Carpenter, et al., Stan: A Probabilistic Programming Language. Journal of Statistical Software 76 (2017).

26. M. Stephens, False discovery rates: a new deal. Biostatistics 18, 275–294 (2017).

27. J. Berkhof, I. van Mechelen, H. Hoijtink, Posterior predictive checks: Principles and discussion. Comput. Stat. 15, 337–354 (2000).

28. J. K. Kruschke, Posterior predictive checks can and should be Bayesian: comment on Gelman and Shalizi, “Philosophy and the practice of Bayesian statistics.” Br. J. Math. Stat. Psychol. 66, 45–56 (2013).

29. A. Gelman, et al., Bayesian Data Analysis, Third Edition (CRC Press, 2013).

30. S. Freytag, L. Tian, I. Lönnstedt, M. Ng, M. Bahlo, Comparison of clustering tools in R for medium-sized 10x Genomics single-cell RNA-sequencing data. F1000Research 7, 1297 (2018).

31. J. Ding, et al., Systematic comparison of single-cell and single-nucleus RNA-sequencing methods. Nat. Biotechnol. 38, 737–746 (2020).

32. T. T. Karagiannis, et al., Single cell transcriptomics reveals opioid usage evokes widespread suppression of antiviral gene program. Nat. Commun. 11, 2611 (2020).

33. Y. Cai, et al., Single-cell transcriptomics of blood reveals a natural killer cell subset depletion in tuberculosis. EBioMedicine 53, 102686 (2020).

34. S. H. Wu, R. S. Schwartz, D. J. Winter, D. F. Conrad, R. A. Cartwright, Estimating error models for whole genome sequencing using mixtures of Dirichlet-multinomial distributions. Bioinformatics 33, 2322–2329 (2017).

35. Z. Dai, S. H. Wong, J. Yu, Y. Wei, Batch effects correction for microbiome data with Dirichlet-multinomial regression. Bioinformatics 35, 807–814 (2019).

36. J. G. Harrison, W. J. Calder, V. Shastry, C. A. Buerkle, Dirichlet–multinomial modelling outperforms alternatives for analysis of microbiome and other ecological count data. Mol. Ecol. Resour. 20, 481–497 (2020).

37. G. Wang, Bayesian and frequentist approaches to multinomial count models in ecology. Ecol. Inform. 61, 101209 (2021).

38. N. Bouguila, Clustering of Count Data Using Generalized Dirichlet Multinomial Distributions. IEEE Trans. Knowl. Data Eng. 20, 462–474 (2008).

39. A. Mishra, C. L. Müller, Robust regression with compositional covariates. Comput. Stat. Data Anal. 165, 107315 (2022).

40. A. Ongaro, C. Cattaneo, Discrete random probability measures: a general framework for nonparametric Bayesian inference. Stat. Probab. Lett. 67, 33–45 (2004).

41. J. Sethuraman, A CONSTRUCTIVE DEFINITION OF DIRICHLET PRIORS. Stat. Sin. 4, 639–650 (1994).

42. S. Mangiola, E. A. Thomas, M. Modrák, A. Vehtari, A. T. Papenfuss, Probabilistic outlier identification for RNA sequencing generalized linear models. NAR Genom Bioinform 3, lqab005 (2021).

43. A. Butler, P. Hoffman, P. Smibert, E. Papalexi, R. Satija, Integrating single-cell transcriptomic data across different conditions, technologies, and species. Nat. Biotechnol. 36, 411–420 (2018).

44. R. A. Amezquita, et al., Orchestrating single-cell analysis with Bioconductor. Nat. Methods 17, 137–145 (2020).

45. M. Trussart, et al., Removing unwanted variation with CytofRUV to integrate multiple CyTOF datasets. Elife 9 (2020).

46. R. P. Schuyler, et al., Minimizing Batch Effects in Mass Cytometry Data. Front.Immunol. 10, 2367 (2019).

47. B. Bodenmiller, et al., Multiplexed mass cytometry profiling of cellular states perturbed by small-molecule regulators. Nat. Biotechnol. 30, 858–867 (2012).

48. S. Van Gassen, B. Gaudilliere, M. S. Angst, Y. Saeys, N. Aghaeepour, CytoNorm: A Normalization Algorithm for Cytometry Data. Cytometry A 97, 268–278 (2020).

49. F. J. Hartmann, et al., Comprehensive Immune Monitoring of Clinical Trials to Advance Human Immunotherapy. Cell Rep. 28, 819–831.e4 (2019).

50. L. Rodriguez, et al., Systems-Level Immunomonitoring from Acute to Recovery Phase of Severe COVID-19. Cell Rep Med 1, 100078 (2020).

51. F. H. Karlsson, et al., Gut metagenome in European women with normal, impaired and diabetic glucose control. Nature 498, 99–103 (2013).

52. D. Gevers, et al., The treatment-naive microbiome in new-onset Crohn’s disease. Cell Host Microbe 15, 382–392 (2014).

53. L. A. David, et al., Diet rapidly and reproducibly alters the human gut microbiome. Nature 505, 559–563 (2014).

54. N. A. Bokulich, et al., Antibiotics, birth mode, and diet shape microbiome maturation during early life. Sci. Transl. Med. 8, 343ra82 (2016).

55. S. J. Song, et al., Cohabiting family members share microbiota with one another and with their dogs. Elife 2, e00458 (2013).

56. M. Pop, et al., Diarrhea in young children from low-income countries leads to large-scale alterations in intestinal microbiota composition. Genome Biol. 15, R76 (2014).

57. P. J. Huber, E. M. Ronchetti, Robust statistics john wiley & sons. New York 1 (1981).

58. C. Jennison, F. R. Hampel, E. M. Ronchetti, P. J. Rousseeuw, W. A. Stahel, Robust Statistics: The Approach Based on Influence Functions. Journal of the Royal Statistical Society. Series A (General) 150, 281 (1987).

59. R. A. Becker, J. M. Chambers, A. R. Wilks, The new S language. Pacific Grove, Ca.: Wadsworth & Brooks, 1988 (1988) (February 25, 2018).

60. H. Wickham, et al., Welcome to the Tidyverse. Journal of Open Source Software 4, 1686 (2019).

61. S. Mangiola, M. A. Doyle, A. T. Papenfuss, Interfacing Seurat with the R tidy universe. Bioinformatics (2021) https://doi.org/10.1093/bioinformatics/btab404.

62. S. Mangiola, R. Molania, R. Dong, M. A. Doyle, A. T. Papenfuss, tidybulk: an R tidy framework for modular transcriptomic data analysis. Genome Biol. 22, 42 (2021).

63. M. Albrecht, P. Donnelly, P. Bui, D. Thain, Makeflow: a portable abstraction for data intensive computing on clusters, clouds, and grids in Proceedings of the 1st ACM SIGMOD Workshop on Scalable Workflow Execution Engines and Technologies, SWEET’ 12., (Association for Computing Machinery, 2012), pp. 1–13.

64. A. Gelman, und Jennifer Hill. 2007. Data analysis using regression and multilevel/hierarchical models.

65. B. Schloerke, et al., GGally: extension to “ggplot2”. R package version 1.4. 0. R Foundation for Statistical Computing (2018).

66. S. Chevrier, et al., A distinct innate immune signature marks progression from mild to severe COVID-19. Cell Rep Med 2, 100166 (2021).

